# Dynamic hinge-motion of PfRIPR revealed by malaria invasion inhibitory antibodies

**DOI:** 10.64898/2026.01.14.699498

**Authors:** Brendan Farrell, Andrew J.R. Cooper, Egle Butkeviciute, Lawrence T. Wang, Emily Egerton-Warburton, Joshua Tan, Matthew K. Higgins

## Abstract

The PfPCRCR complex is essential for invasion of human erythrocytes by the deadliest malaria parasite, *Plasmodium falciparum*. Antibodies against each subunit of PfPCRCR prevent erythrocyte invasion and the PfRH5 component is currently the most advanced blood-stage malaria vaccine. Central within PfPCRCR is PfRIPR. This complex molecule contains a core and a flexible tail and allows PfPCRCR to bridge the parasite and erythrocyte during invasion. In this study, we generated a small panel of human monoclonal antibodies against PfRIPR. We structurally characterised four PfRIPR tail-binding antibodies in complex with PfRIPR fragments. We show that growth-inhibitory antibody RP.012 induces a kink in the PfRIPR tail while non-inhibitory antibodies do not. Furthermore, we show that these four antibodies modulate each other, either through antagonism or by acting synergistically. These studies have implications for the design of PfRIPR-based vaccine immunogens and indicate that the tail of PfRIPR undergoes essential conformational changes during erythrocyte invasion.

## Introduction

Invasion of erythrocytes by *Plasmodium falciparum* is an active multi-stage process orchestrated by a repertoire of parasite surface proteins^1^. Among these, the PfPCRCR complex forms an essential bridge between parasite and erythrocyte membranes^2,3^. PfPCRCR contains five components (PfPTRAMP, PfCSS, PfRIPR, PfCyRPA and PfRH5) (Figure 1A), of which PfRH5 has emerged as the leading candidate for a blood-stage malaria vaccine^1^. The interaction between PfRH5 and its erythrocyte receptor basigin is essential for invasion^4-6^ and allows the parasite-associated PfPCRCR complex to bind the erythrocyte membrane^2,3^.

**Figure 1:**
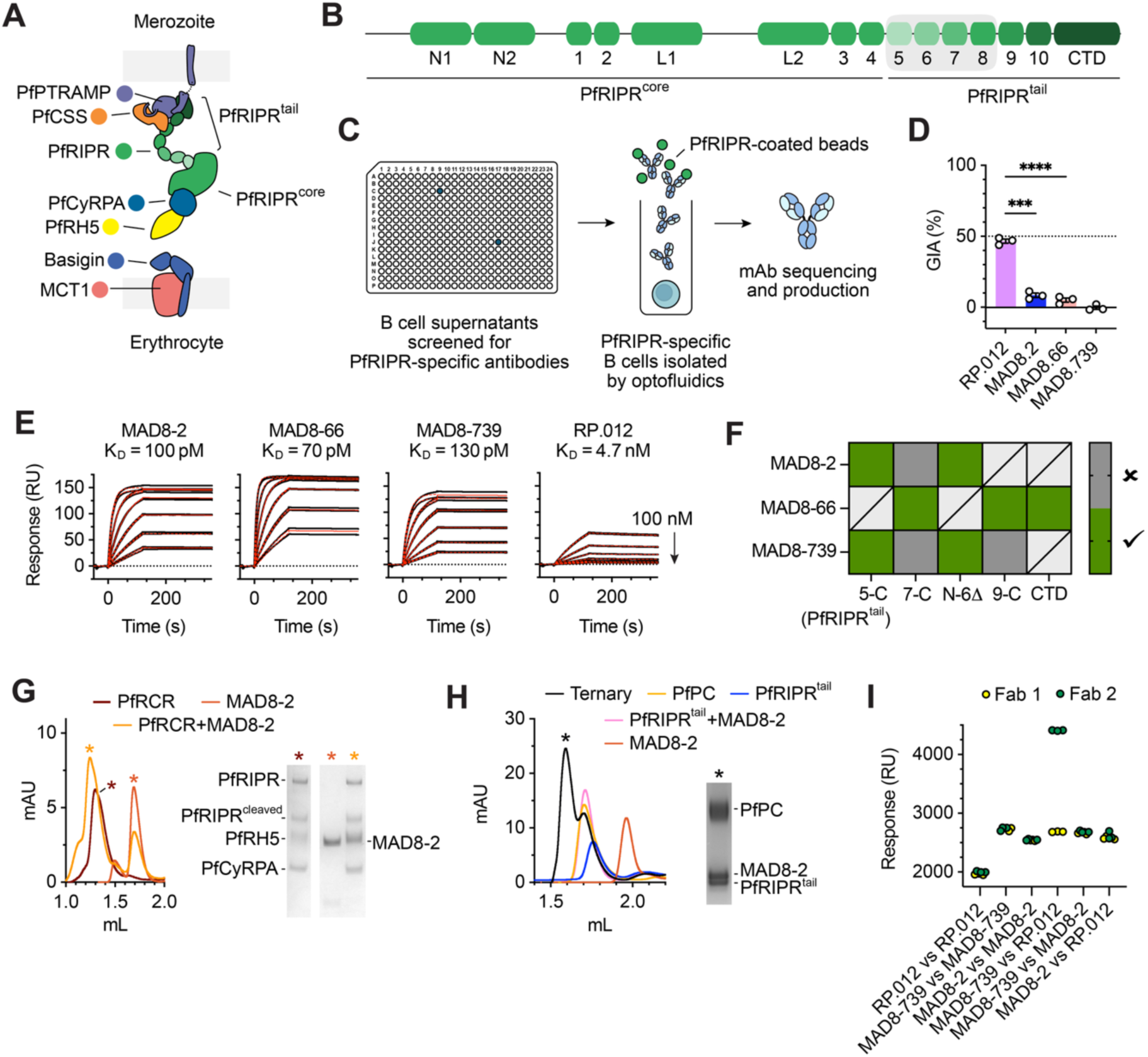
Isolation and characterisation of PfRIPR antibodies. (A) Illustration of the PfPCRCR complex which forms a bridge between the merozoite and erythrocyte membranes upon binding of basigin on the erythrocyte surface^2,3^. Note that the precise tethering arrangement of the PfPCRCR complex via the transmembrane domain of PfPTRAMP is unknown (indicated by a dashed line). (B) Schematic of the domain composition of PfRIPR with the location of its known neutralising epitopes in EGF-like domains 5-8 highlighted in grey^8,16-18^. (C) Pipeline to isolate human PfRIPR antibodies from malaria-exposed individuals. (D) Single-point GIA analysis of each PfRIPR monoclonal antibody assayed at 1 mg/mL. The GIA of RP.012 is significantly greater than those of MAD8-2 and MAD8-66 (P=0.0001 and P<0.0001); n=3, Brown-Forsythe and Welch ANOVA test with Dunnett’s T3 multiple comparisons test; individual values (circles), mean (bar height) and standard deviation shown. (E) SPR traces (black) showing binding of PfRIPR to each monoclonal antibody, their kinetic fits (red), and affinity K_D_ values; n=4, 2 repeats shown. (F) Epitope binning of PfRIPR antibodies to fragments of PfRIPR. Green bins indicate that binding was observed between Fab fragments and PfRIPR fragments, while dark grey bins indicate that no binding was observed. Combinations that were not tested are crossed through. Binning analyses are shown in Figure S2A-D. (G) Size exclusion chromatography (SEC) traces of MAD8-2 Fab fragment (red), the PfRCR complex (brown), and their mixture (orange). Peak fractions (as indicated by asterisks) were analysed by SDS-PAGE confirming co-elution of all components. Equivalent analyses for MAD8-66, MAD8-739 and RP.012 are shown in Figure S2E. (H) SEC traces of MAD8-2 Fab fragment (red), PfRIPR tail (blue), PfPTRAMP-PfCSS (PfPC, yellow), PfRIPR tail plus MAD8-2 (pink), and the complex of all components (black). The peak fraction of this complex (indicated by an asterisk) was analysed by SDS-PAGE confirming co-elution of all components. Equivalent analyses for MAD8-66, MAD8-739 and RP.012 are shown in Figure S2F. (I) Competition binning of EGF5-8 binding antibodies by SPR. The binding response (RU) to PfRIPR following the injection of the first Fab fragment of an antibody pair (yellow circles), then the response after the sequential injection of the second Fab fragment of the pair after prior binding of the first (green circles); individual data points shown, n=3.

A vaccine based on full-length PfRH5, known as RH5.1, has advanced through clinical trials and has an efficacy of ∼55% against clinical malaria when administered to infants in a seasonal malaria setting in Burkina Faso^7^. This shows the promise of PfRH5 as a blood-stage vaccine but also emphasizes the need for further improvement. Structure-guided approaches are therefore currently being used to generate next-generation PfRH5-based vaccines^1^. In addition, as each component of PfPCRCR is individually essential for invasion^3,8,9^ and each raises invasion-inhibitory antibodies or nanobodies^3,5,8,10-20^, they are all potential vaccine candidates, alone or in combination with PfRH5.

PfRIPR is the largest and least understood component of PfPCRCR. It has a complex multidomain architecture consisting of two distinct parts: a core of 8 domains which fold into a compact structure and a flexible tail formed from six EGF-like domains and a C-terminal domain^2^ (Figure 1A-B). PfRIPR is central to the PfPCRCR complex, interacting with PfCyRPA-PfRH5 through its core^2,21^ and with the PfPTRAMP-PfCSS heterodimer through its tail^2,22^. PfPTRAMP contains a transmembrane helix^23^ which immobilises PfPCRCR to the merozoite surface, such that the interaction of PfRH5 with basigin completes a bridge between merozoite and erythrocyte^2,3^. However, whether PfRIPR is simply a bridge, or plays a more active role in invasion, is not known.

While antibody-mediated responses to PfRH5 have been extensively studied^5,10-12,24^, little is known about how PfRIPR-targeting antibodies inhibit invasion. There is no consistent correlation between seropositivity against PfRIPR and protection from clinical malaria in endemic regions^16,25,26^, and polyclonal sera raised by immunisation with full-length PfRIPR has low growth-inhibitory activity^18^. However, EGF-like domains 5-8 in the tail of PfRIPR can elicit polyclonal antibodies with better growth-inhibitory activity than those elicited using full-length PfRIPR and the epitopes for growth inhibitory antibodies raised through PfRIPR immunisation have been mapped by depletion studies to this region^16-18^ (Figure 1B). Indeed, while few antibodies against PfRIPR have been isolated, those with growth-inhibitory activity, including RP.012, bind epitopes within EGF5-8^16,18^. These findings led to the design of the R78C vaccine, which contains EGF-like domains 7 and 8 of PfRIPR fused to PfCyRPA, provided in combination with RH5.1^18^. Preclinical studies indicated that IgG induced by R78C showed a small but significant increase in growth-inhibitory activity when compared with those induced with PfRH5.1, and R78C is now in a Phase I clinical trial (NCT05385471)^18^.

Despite the early adoption of a specific region of PfRIPR in a vaccine, there is still a lot to learn about how PfRIPR functions during invasion or how it is inhibited by antibodies. Few monoclonal antibodies (mAbs) have been identified and, with no structures of mAbs bound to PfRIPR, we do not understand how they block invasion, limiting our ability to rationally design the most effective vaccine immunogen based on PfRIPR. Here, we isolated and characterised a small panel of PfRIPR antibodies acquired naturally by individuals in a malaria-endemic region of Mali, which we combine with the inhibitory antibody RP.012. Studies of this panel reveals that PfRIPR has a dynamic role which is exploited by growth-neutralising antibodies. Moreover, we find that antibodies binding to spatially distant sites in PfRIPR synergise with one another, suggesting the inclusion of discontinuous regions of PfRIPR in future vaccine design.

## Results

### Isolation of a panel of human PfRIPR-targeting antibodies

We isolated a panel of PfRIPR-specific monoclonal antibodies (mAbs) from a cohort of 758 individuals living in Kalifabougou, Mali. This is a highly endemic region for *P. falciparum*, with an estimated two infectious bites per person per day at the peak of the malaria season. Since to our knowledge there are no ongoing or planned clinical trials for vaccination with full-length PfRIPR, natural infection presents an invaluable source of human PfRIPR-specific mAbs. From this cohort, we isolated the first human anti-PfRIPR mAbs to date. IgG^+^ memory B cells from malaria-exposed individuals were screened sequentially using flow cytometry and an optofluidics platform to identify cells that secrete PfRIPR-binding antibodies. B cells of interest were extracted and sequenced, and corresponding mAbs were recombinantly expressed and tested to confirm binding to PfRIPR (Figures 1C and S1A). These are among the first mAbs isolated as part of a larger-scale discovery effort to characterize the human immune response to PfRIPR.

From this small pilot panel of human antibodies, we selected three IgG1 antibodies for further analysis: MAD8-2, MAD8-66 and MAD8-739 (Figure S1). Together with the previously identified growth-inhibitory murine antibody RP.012^18^, which we recombinantly expressed as a chimeric human IgG1 antibody, these formed our panel. We measured the growth inhibition activity (GIA) of each antibody, finding that only RP.012 reduced erythrocyte invasion, with a GIA of ∼46% at 1 mg/mL consistent with published data^18^ (Figure 1D). None of our three naturally-acquired human mAbs showed GIA greater than ∼8% at 1 mg/mL (Figure 1D). We next determined the binding affinity of each mAb for PfRIPR using surface plasmon resonance (SPR) (Figure 1E; Table S1). Each human mAb bound PfRIPR with sub-nanomolar affinity (∼70-130 pM), indicating that their poor neutralising capacity is not due to poor affinity for PfRIPR. RP.012 also bound tightly (∼5 nM), but with a slower association rate.

The epitope of RP.012 has already been mapped to EGF-like domains 6-7^18^. We next assessed binding of Fab fragments of our human mAbs to recombinant PfRIPR fragments. Both MAD8-2 and MAD8-739 bound fragments 5-C (PfRIPR^tail^, comprising EGF5 to the C-terminus of PfRIPR) and N-6Δ (N-terminus to EGF6 except the L1-L2 loop) but not PfRIPR fragment 7-C (EGF7 to the C-terminus), indicating that they each have an epitope within EGF-like domains 5-6 (Figures 1F and S2A-B). This was unexpected given that all previously characterised mAbs that bind to EGF5-8 are growth-inhibitory^16,18^. Conversely, MAD8-66 bound each of 7-C, 9-C (EGF9 to the C-terminus), and the C-terminal domain (CTD), indicating an epitope within the CTD (Figures 1F and S2C-D).

Next, to determine whether each epitope is accessible in the PfPCRCR complex, we assembled complexes of these Fab fragments with PfRCR or PfPTRAMP-PfCSS-PfRIPR (PfPCR). All four antibodies co-eluted in complex with PfRCR, indicating that they do not disrupt binding of PfRIPR to PfRH5 or PfCyRPA (Figures 1G and S2E). To assess whether these antibodies inhibited binding of PfPTRAMP-PfCSS to PfRIPR, we focused on PfRIPR tail, as all four mAbs and PfPTRAMP-PfCSS bind within this fragment^2,22^. All four antibodies co-eluted with the PfPTRAMP-PfCSS-PfRIPR^tail^ complex at an earlier elution volume than any component of the complex individually or the Fab-PfRIPR^tail^ complex, as exemplified for MAD8-2, demonstrating that they do not disrupt the interaction of PfRIPR with the PfPTRAMP-PfCSS heterodimer (Figures 1H and S2F). While the elution profiles of the MAD8-2, MAD8-66 and MAD8-739 complexes were similar, the RP.012 complex eluted at a slightly later elution volume for both PfRCR and PfPCR, which may indicate that RP.012 induces a conformational change in PfRIPR (Figures S2E-F).

Finally, as three of our antibodies (RP.012, MAD8-2 and MAD8-739) bind within EGF-like domains 5-7 of PfRIPR, we determined whether they compete with one another by immobilising full-length PfRIPR on an SPR chip and sequentially injecting Fab fragments at saturating concentration (Figures 1I and S2G). RP.012 bound to PfRIPR pre-saturated with MAD8-739, but MAD8-2 did not. Conversely, PfRIPR pre-saturated with MAD8-2 did not bind RP.012. Therefore MAD8-2 shares an overlapping epitope with both MAD8-739 and RP.012, but MAD8-739 and RP.012 do not compete with one another.

In summary, our panel consists of four antibodies that interact with exposed epitopes within the PfPCRCR complex, with two binding to EGF5-6 (MAD8-2 and MAD8-739), one binding to EGF6-7 (RP.012), and one binding to the CTD of PfRIPR (MAD8-66). Only RP.012 exhibits GIA activity, despite our human mAbs binding with high affinity, and two of these antibodies (MAD8-2 and MAD8-739) interacting with EGF5-8, reported to contain all neutralising epitopes of PfRIPR^18^. Moreover, one of these (MAD8-2) directly competes with RP.012. This indicates that binding to PfRIPR EGF5-8 alone is not sufficient to impede erythrocyte invasion and that specific surfaces must be engaged. It also shows that non-inhibitory antibodies targeting this region of PfRIPR can interfere with the function of growth-inhibitory antibodies, showing the need for caution in vaccine design.

### Growth-inhibitory antibody RP.012 bends PfRIPR

To investigate why RP.012 is uniquely GIA active among the three EGF5-8 binders in this panel, we sought to determine the structure of each antibody complex using X-ray crystallography. While this was unsuccessful for MAD8-2, we obtained crystal structures of PfRIPR EGF6-7 bound to the Fab fragment of RP.012 at 2.20 Å resolution (Figure 2A; Table S2), and of PfRIPR EGF5-6 and the single chain variable fragment (scFv) of MAD8-739 at 2.80 Å resolution (Figure 2B; Table S2).

**Figure 2:**
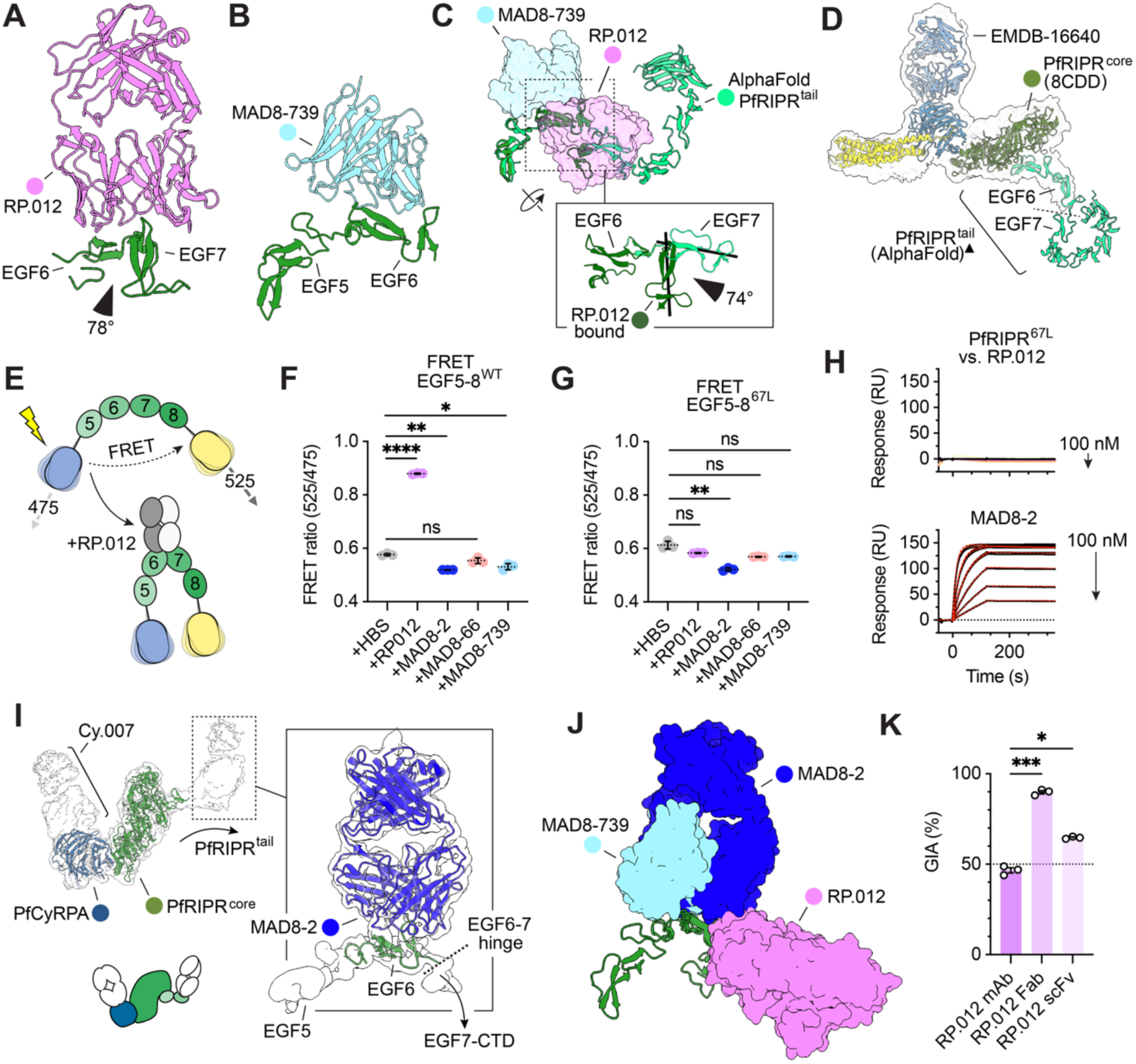
Structural and functional characterisation of PfRIPR EGF5-8 binding antibodies. (A) Crystal structure of RP.012 Fab fragment (pink) and PfRIPR EGF6-7 (green) shown in cartoon representation, with the angle between these domains indicated. (B) Crystal structure of MAD8-739 scFv fragment (light blue) and PfRIPR EGF5-6 (green). (C) Alignment of the RP.012-bound structure, MAD8-739-bound structure and AlphaFold prediction of PfRIPR^tail^. PfRIPR in each of these is shown in cartoon representation with RP.012 and MAD8-739 shown as transparent surfaces. The inset shows a comparison of the position of EGF6 and EGF7 in the RP.012 bound state (dark green) and the AlphaFold prediction of PfRIPR^tail^ (light green). Black sticks show the centre of mass of EGF7 in each of these models and the angle between them is indicated. (D) Cryo-EM volume EMDB-16640 (transparent surface) and a docked composite model of Cy.003-bound PfRCR comprised of PDB entry 8CDD and the AlphaFold prediction of PfRIPR^tail^ (▲)^2^, illustrating a sudden decrease of cryo-EM density approximately between EGF6 and 7. (E) Design of a PfRIPR FRET probe comprising PfRIPR EGF5-8 flanked by mCerulean (blue) and mCitrine (yellow) which emit fluorescence at 475 nm and 525 nm respectively^29^. Excitation of the FRET probe by illumination at 420 nm (lightning bolt) causes FRET transfer between these probes (dashed arrow). Based on the crystal structure, RP.012 is expected to induce a conformational change bringing these fluorescent proteins closer to one another as illustrated, which would result in an increase in FRET efficiency. (F) FRET efficiency (quantified as the FRET ratio between emission at 525 and 475 nm) of the wild-type PfRIPR FRET probe (FRET EGF5-8^WT^) in the presence of HBS, or Fab fragments of EGF5-8 binding antibodies RP.012, MAD8-2, or MAD8-739, or the CTD-binding antibody MAD8-66. RP.012 induces an increase in FRET efficiency (P<0.0001), while FRET efficiency in the presence of MAD8-2 and MAD8-739 is slightly reduced (P=0.0047 and P=0.0220). Non-binder MAD8-66 did not induce a significant change (P=0.1207); n=3 technical replicates, Brown-Forsythe and Welch ANOVA test with Dunnett’s T3 multiple comparisons test; individual values (circles), mean (dashed line) and standard deviation are shown. (G) FRET efficiency of the EGF6-7 cysteine-locked variant of the FRET probe (FRET EGF5-8^67L^) in the presence of HBS or Fab fragments as in Figure 2F, demonstrating that RP.012, MAD8-739 and non-binder MAD8-66 do not induce a change in FRET efficiency with respect to HBS (P=0.1504, P=0.0931 and P=0.0883). The reduction in FRET efficiency in the presence of MAD8-2 was significant (P=0.0063); n=3 technical replicates, Brown-Forsythe and Welch ANOVA test with Dunnett’s T3 multiple comparisons test; individual values (circles), mean (dashed line) and standard deviation are shown. (H) Representative SPR traces measuring the interaction of EGF6-7 cysteine-locked PfRIPR (PfRIPR^67L^) with RP.012 or MAD8-2 and their kinetic fits for MAD8-2 (red). RP.012 does not interact with PfRIPR^67L^ (n=3, 2 shown), while MAD8-2 does (n=4, 2 shown). (I) Cryo-EM volume of MAD8.2-bound PfRIPR-PfCyRPA-Cy.007 (transparent surface) and schematic below showing that the MAD8-2 Fab fragment binds at the end of density corresponding to the tail of PfRIPR. Shown inset, cryo-EM volume of MAD8-2 Fab fragment and PfRIPR and model of MAD8-2 (dark blue) and PfRIPR EGF6 (green) shown in cartoon representation. The location of the hinge between EGF-like domains 6 and 7 is shown by a dotted line. (J) Alignment of structures of RP.012- (pink), MAD8-2- (dark blue) and MAD8-739-bound PfRIPR (light blue), demonstrating steric clashes of MAD8-2 with both MAD8-739 and RP.012. Each antibody is shown in surface representation and PfRIPR in cartoon representation. (K) Single-point GIA of RP.012 as a monoclonal antibody (mAb), a Fab fragment, and an scFv fragment demonstrating that smaller fragments of RP.012 also have GIA activity, at a level significantly greater than RP.012 mAb (P=0.0002 and P=0.0106); n=3, Brown-Forsythe and Welch ANOVA test with Dunnett’s T3 multiple comparisons test; individual values (circles), mean (bar height) and standard deviation are shown. RP.012 mAb was assayed at ∼6.9 µM (corresponding to 1 mg/mL) and its Fab and scFv fragments at ∼13.8 µM, ensuring an equal number of paratopes.

RP.012 binds at the interface between EGF-like domains 6 and 7, such that lines through the two domains lie at 78° to one another (Figure 2A). The RIPR-RP.012 interface has a buried surface area of 827 Å^2^ and is driven by interactions involving CDRs H1, H3, L1-3 and some light and heavy chain framework regions (Table S3). In contrast, the epitope of MAD8-739 is contained mostly within EGF6 of PfRIPR with a buried surface area of 789 Å^2^ (Figure 2B). While EGF6 has a canonical EGF-like domain fold comprising a major sheet (β1-β2) and a minor sheet (β3-β4), containing three disulphide bonds (Figure S3A), EGF5 of PfRIPR has a non-canonical fold containing an additional segment that is similar to a short beta sheet (β5-β6) held in place by a further disulphide bond, similar to EGF-like domains of laminins^27^ (Figure S3A). MAD8-739 interacts with this extra element (β5-β6) of EGF5 and the major sheet of EGF6 through CDRs H1-3, L1 and L3 (Table S4).

Previous AlphaFold^28^ predictions suggest PfRIPR tail to form an approximately linear array of domains^2^ (Figure S3B). Structural alignment of this prediction with our RP.012-bound structure reveals a bend between EGF6 and 7 that rotates EGF7 by 74° (Figure 2C). In cryo-EM data of the PfRCR complex (EMDB-16640)^2^, a sudden decrease in map density is seen at the approximate location of EGF6 and 7, which was previously attributed to conformational flexibility of the tail^2^ (Figure 2D). The bend identified in the presence of RP.012 is therefore consistent with the mAb capturing and exploiting flexibility of the PfRIPR tail to induce a specific conformation. In contrast, the MAD8-739-bound structure of EGF5-6 is similar to the AlphaFold model (Figure 2C), suggesting that MAD8-739 does not bend PfRIPR. This raises the question of whether the growth-inhibitory activity of RP.012 correlates with its ability to stabilise or induce a bent conformation of the PfRIPR tail.

To assess whether RP.012 can indeed stabilise a bent conformation of PfRIPR in solution, we designed an *in vitro* Förster resonance energy transfer (FRET) probe containing PfRIPR EGF5-8 flanked by mCerulean and mCitrine^29^ (Figure 2E), then assessed the effect of each EGF5-8 binding antibody on its FRET efficiency, which is influenced by fluorophore separation^30^. MAD8-66 (the CTD-binding antibody) was used as a negative control to assess for molecular crowding effects. RP.012 increased FRET efficiency of the probe significantly (Figure 2F), while MAD8-2 and MAD8-739 decreased FRET efficiency (Figure 2F). This is consistent with RP.012 but not MAD8-2 nor MAD8-739 stabilising a bent conformation of the PfRIPR tail. We next designed a disulphide lock between EGF6 and 7 at residues H804C and F835C to hold PfRIPR^tail^ in an extended conformation (Figure S3C) and verified this by iodoacetamide labelling and intact mass spectrometry analysis of PfRIPR EGF5-8 with this designed disulphide (EGF5-8^67L^) (Figure S3D). RP.012 did not increase FRET efficiency of a probe containing EGF5-8^67L^ (Figure 2G). Moreover, unlike MAD8-2, RP.012 could not bind PfRIPR^67L^, indicating that its epitope is no longer accessible in this conformationally-restrained version of PfRIPR (Figure 2H; Table S5).

As MAD8-2 bound to PfRIPR did not crystallise, we determined its structure using single particle cryo-EM. Given the low molecular weight of PfRIPR EGF5-6 bound to MAD8-2 (only ∼60 kDa with a Fab fragment), we reconstituted PfRCR with Fab fragments of MAD8-2 and the PfCyRPA antibody Cy.007^15^. We used the cysteine-locked version of PfRIPR in this complex, aiming to visualise the remainder of the tail of PfRIPR that is missing in prior studies^2^ (Figure S4A). We obtained a volume for this MAD8-2-PfRIPR-PfCyRPA-Cy.007 complex at a global 3.0 Å resolution that showed MAD8-2 bound at the tip of density corresponding to EGF5 and 6 of the tail of PfRIPR (Figure 2I; Table S6). MAD8-2 was not well-resolved in this map and, while further density for the tail of PfRIPR was visible in 2D class averages, this was not resolved in 3D (Figure S4B). Separately, we obtained an additional map of a MAD8-2-bound PfRIPR fragment at a nominal 3.5 Å resolution (Table S6; Figure S4B), into which we could unambiguously dock AlphaFold3 models of MAD8-2 variable domain and PfRIPR EGF6 (Figure 2I). While this map is of insufficient quality to confidently define residue-level interactions, it shows that MAD8-2 binds entirely within EGF6 using each CDR except CDR L2. Alignment of our PfRIPR-antibody structures confirms that MAD8-2 and MAD8-739 share overlapping epitopes, but that MAD8-739 and RP.012 do not (Figure 2J). MAD8-2 binds further along EGF6 than MAD8-739, causing it to share some binding surface with RP.012 (Figure S3E).

Combining these structures, FRET data, and kinetic binding analysis suggests a model in which RP.012 induces or captures a bent conformation of the PfRIPR tail, at a location that cryo-EM data^2^ indicates is flexible. If RP.012 exerts its growth-inhibitory activity through this mechanism, we would expect smaller antibody fragments of RP.012 to also induce this conformational change and be growth-inhibitory. To test this, we conducted single point GIA assays for the mAb, Fab fragment and scFv forms of RP.012, using molar equivalents of the paratope (Figure 2K). These were all growth-inhibitory, with the Fab and scFv fragments outperforming the mAb with ∼90% and ∼65% inhibition (Figure 2K), consistent with the conformational change induced or stabilised by RP.012 being responsible for its growth-inhibitory activity.

### Building towards a more complete structural model of PfRIPR

Our cryo-EM volume of MAD8-2-bound PfRIPR-PfCyRPA-Cy.007 allowed us to build a structure of PfRIPR spanning from the N-terminus to the end of EGF5 (residues 25-759 except disordered loops) (Figure 3A; Table S6). This extends the known structure of PfRIPR by 9 and 53 residues at the N and C-termini respectively and shows that EGF5 contacts the underside of the PfRIPR core, beneath domains L1 and L2, making contacts through both its major (β1-β2) and minor sheets (β3-β4) with L1, L2, and the N-terminus of PfRIPR (Figure 3A). This agrees with prior crosslinking data of the PfRCR complex, which identified a crosslink between EGF5 and L1^2^. The N-terminus of PfRIPR interacts with domains EGF5, L1 and L2. This interaction network determines the angle at which the tail of PfRIPR projects from the core.

**Figure 3:**
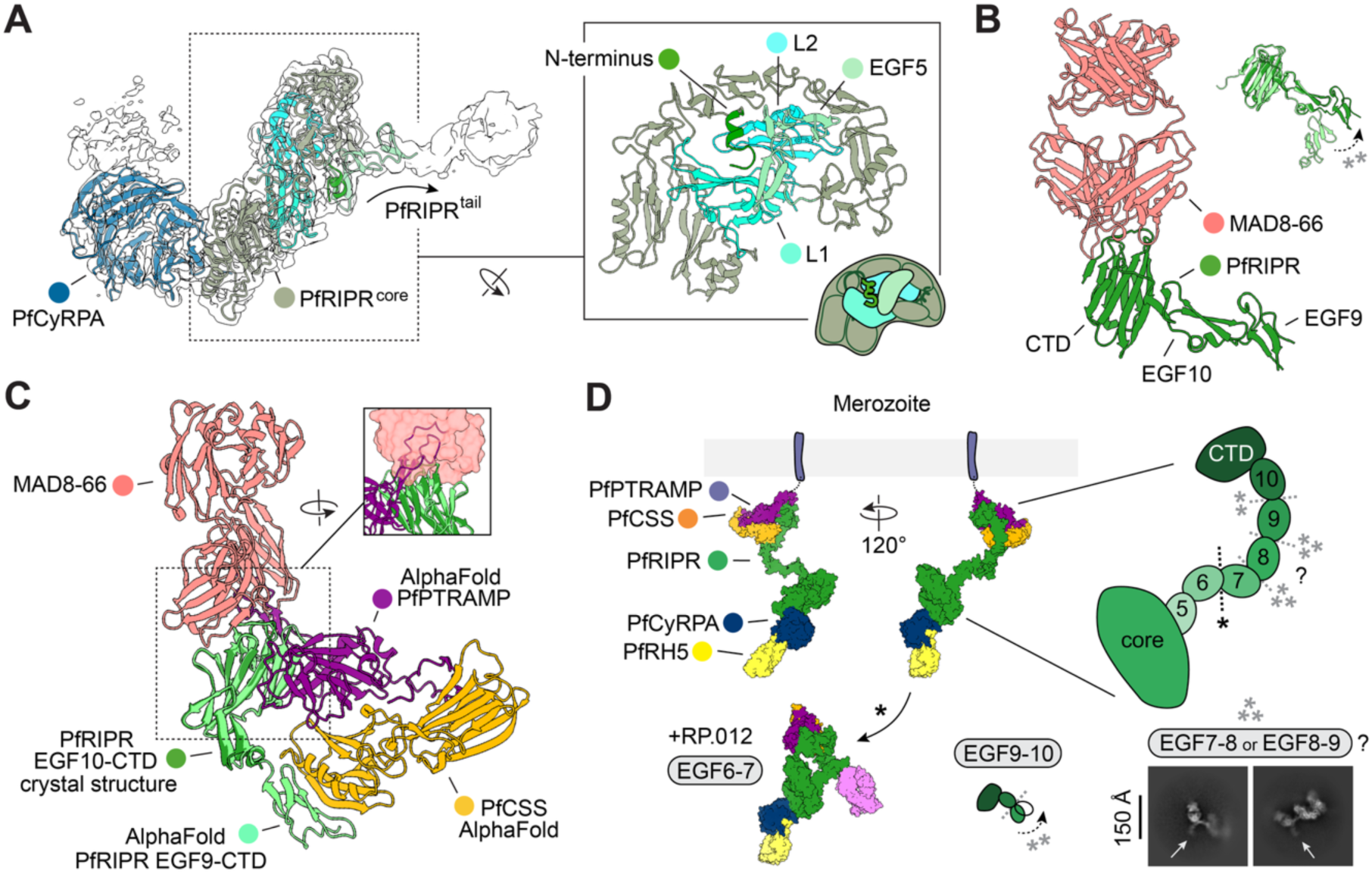
Building a more complete structural understanding of PfRIPR. (A) Cryo-EM volume of MAD8-2-bound PfRIPR-PfCyRPA with a new model of PfRIPR comprising the N-terminus to EGF5 shown in cartoon representation. The inset shows that the structure reveals the tail of PfRIPR interacting with PfRIPR^core^ through a platform of interactions comprised of domains L1 (mint), L2 (cyan), EGF5 (light green) and the N-terminus (green), as illustrated schematically. (B) Crystal structure of MAD8-66 (salmon) bound to PfRIPR EGF9-CTD (green). Inset upper right, an alignment of EGF9-CTD from the AlphaFold prediction of PfRIPR^tail^ (light green) and the MAD8-66-bound structure, illustrating shifting of EGF9. (C) Alignment of the crystal structure of MAD8-66-bound PfRIPR (truncated to remove EGF9 for clarity,) and the AlphaFold prediction of PfPTRAMP (purple), PfCSS (orange) and PfRIPR^tail^ (light green, trimmed back to EGF9 for clarity) complex. The AlphaFold model was additionally trimmed to remove N and C-termini with low pLDDT scores (Figure S5C, residues L42-N50 and N297-D352 of PfPTRAMP, and residues Q21-K27 of CSS). Inset upper right, MAD8-66 shown with a transparent pink surface to illustrate a potential steric clash with PfPTRAMP. (D) Composite model of the PfPCRCR complex shown in surface representation, combining experimentally determined structures and the AlphaFold prediction of the PfPTRAMP-PfCSS-PfRIPR^tail^ complex. Inset, an expanded view of PfRIPR with potential sites of flexibility in its tail indicated by dashed lines with asterisks showing supporting experimental evidence. * - RP.012 induces a conformational change between EGF6-7 (Figure 2), ** - crystal contacts induce a conformational change between EGF9-10 (Figures 3B and S5A-B), *** - 2-dimensional cryo-EM classes of MAD8-2-bound PfRIPR^67L^-PfCyRPA-Cy.007 show incomplete tail density (highlighted by white arrows) estimated to drop off approximately between EGF7-8 or EGF8-9.

We next determined the crystal structure of MAD8-66 bound to PfRIPR EGF9-C to a resolution of 2.25 Å (Figure 3B; Table S2). Consistent with epitope binning (Figure 1F), MAD8-66 interacts solely with the CTD of PfRIPR, contacting both beta sheets of its beta-sandwich fold, plus the linker following EGF10. This interaction buries 1020 Å^2^ using all CDR loops (Table S7). Our crystal structure aligns well with the AlphaFold model of the tail of PfRIPR except at EGF9, which both pivots away with respect to the predicted model and undergoes a structural rearrangement of its minor sheet (β3-β4) into a single turn helix (Figures 3B and S5A). As EGF9 makes crystal contacts here, this may be a crystallisation artefact (Figure S5B), but we note that the Predicted Aligned Error (PAE) matrix of the AlphaFold prediction of PfRIPR^tail^ suggests that EGF10-CTD is a structural unit which is distinct from the rest of the tail (Figure S3B). This conformational difference at the interface between EGF9 and 10 may therefore reflect a genuine location of further flexibility in the tail of PfRIPR.

No experimental structure has been resolved for the PfPTRAMP-PfCSS-PfRIPR (PfPCR) complex. To visualise MAD8-66 binding in context of PfPCRCR, we therefore compared our structure of MAD8-66-bound PfRIPR to the AlphaFold3 prediction of PfPCR (Figures 3C and S5C). This alignment suggests that MAD8-66 would sterically clash with the thrombospondin repeat (TSR) domain of PfPTRAMP, which is predicted to interact with the CTD of PfRIPR, despite MAD8-66 co-eluting with the PfPCR complex (Figure S2F). While the PAE matrix of this AlphaFold prediction shows that the predicted positional error of the TSR domain with respect to PfRIPR CTD is low, implying confidence in their predicted interface (Figure S5C), it is notable that PfCSS is reported to drive the interaction of PfPTRAMP-PfCSS heterodimer with PfRIPR *in vitro* and that no direct interaction between PfPTRAMP and PfRIPR has been experimentally demonstrated^3,22^. This casts doubt on the AlphaFold prediction of this complex, which shows an extensive interface between PfPTRAMP and PfRIPR. However, we note that single particle cryo-EM 2D class averages of the equivalent complex in *Plasmodium knowlesi* are similar to projections of its AlphaFold prediction^22^, lending support to the model. One possible explanation to rationalise these data is that the interaction between PfCSS and PfRIPR is sufficient for formation of PfPCR and that this complex remains associated even if MAD8-66 disrupts the interaction between PfPTRAMP and PfRIPR.

To model the intact PfPCRCR complex, we generated a composite model using known structures of PfRCR (PDB entry 8CDD), PfRIPR N-EGF5 and PfRIPR EGF5-6 (described herein), and the AlphaFold predicted structures of the PfRIPR tail and the PfPCR complex (Figure 3D). This provides a static snapshot of PfPCRCR. However, our data indicate that PfRIPR is flexible, notably between EGF6-7 as exploited by RP.012, but also potentially at other locations in the tail of PfRIPR such as between EGF9-10, and most likely between EGF7-8 or EGF8-9, as seen from loss of PfRIPR tail density in cryo-EM 2D class averages of MAD8-2-bound PfRIPR^67L^-PfCyRPA-Cy.007 (Figure 3D). Further composite modelling using our RP.012-bound PfRIPR crystal structure to guide the alignment suggests that RP.012 would induce the PfPCRCR complex to fold onto itself in a more compact state (Figure 3D), perhaps providing a clue to its mode of action.

### PfRIPR-targeting mAbs can act synergistically

Our data show that PfRIPR EGF5-8 is the target of both neutralising and non-neutralising antibodies, with different antibody pairs either competing or binding simultaneously (Figures 1I and S2G). Polyclonal sera most likely contain a mixture of these antibodies. To test whether antibody combinations affect growth-inhibitory activity, we measured GIA of pairwise equimolar combinations of our EGF5-8 binding mAbs in a single-point GIA assay, holding each antibody at 1 mg/mL (Figure 4A). The observed GIA of RP.012 plus MAD8-2 was only ∼16% compared to 46% for RP.012 alone (Figure 1D), indicating that MAD8-2 can indeed compete with and antagonise RP.012. In contrast, the non-competing mAbs MAD8-739 and RP.012 showed a combined GIA of ∼65% inhibition, greater than the calculated Bliss Additivity^31^ for this combination of 46% (Figure 4A). This suggests that these antibodies, which bind simultaneously to sites close together on PfRIPR, may show some weak synergy. The neutralising capacity of a polyclonal pool of EGF5-8 antibodies is therefore likely to be a complex combination of synergy and antagonism.

**Figure 4:**
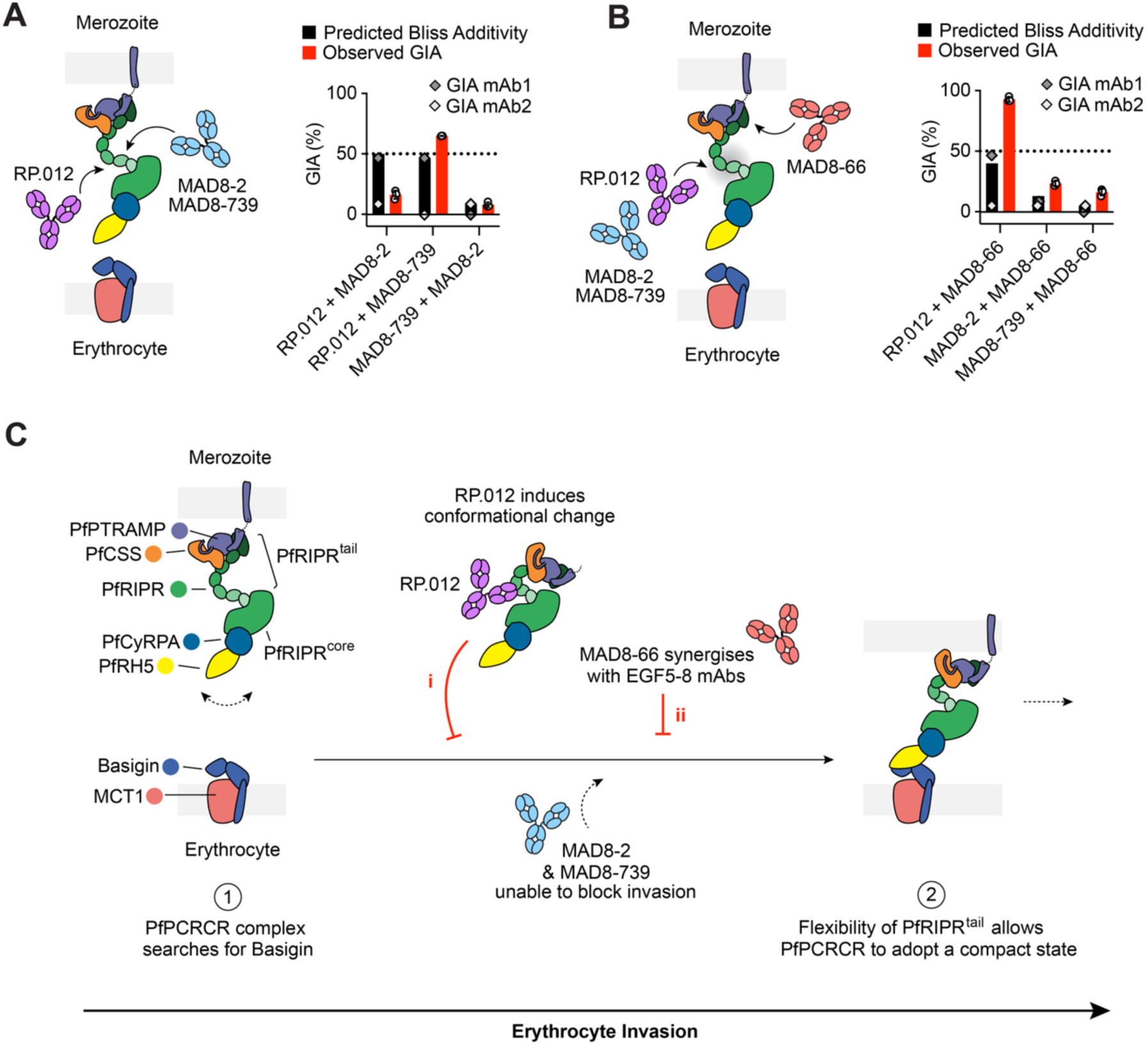
Synergy and competition between PfRIPR antibodies. (A) Schematic diagram (left) and single point GIA (right) of pairwise combinations of PfRIPR EGF5-8 binding antibodies held at 1 mg/mL each, demonstrating both competition and synergy. For each combination, diamonds show the mean GIA of mAb1 (dark grey) and mAb2 (light grey) (as measured in Figure 1D), and a black bar shows the expected Bliss Additivity of these two mAbs. The adjacent red bar shows the mean observed GIA of these combined mAbs with data points (white circles, n=3) and standard deviation. (b) Schematic diagram (left) and single point GIA (right) of pairwise combinations of CTD-binding MAD8-66 and each EGF5-8 binding antibody, each held at 1 mg/mL demonstrating synergy of MAD8-66 with EGF5-8 binding mAbs. Data are presented as in Figure 4A; n=3. (c) Schematic illustrating a model for how the PfPCRCR complex may act during invasion of erythrocytes. Flexibility in the tail of PfRIPR may allow the PfPCRCR complex to search for basigin on the erythrocyte surface. On binding to basigin, we propose that the PfPCRCR complex must undergo a conformational change to allow the parasite and erythrocyte membranes to come closer together, to allow unknown downstream events that lead to rhoptry discharge. PfRIPR antibodies may impede invasion in several ways, for example by (i) GIA-positive mAbs such as RP.012 inducing conformational changes that prevent basigin binding or are incompatible with invasion, or (ii) by synergy between mAbs which bind the CTD and EGF5-8 preventing the required conformational changes.

Polyclonal responses will also contain antibodies targeting elsewhere in PfRIPR. We therefore measured GIA for each of our EGF5-8 antibodies combined with MAD8-66, our CTD-binding antibody. Unexpectedly, we found that MAD8-66 can synergise with EGF5-8 antibodies (Figure 4B). MAD8-66 and RP.012 had a combined GIA of ∼93%, substantially improving the neutralising capacity of RP.012. Moreover, MAD8-66 when combined with MAD8-2 or MAD8-739 showed GIA of ∼23% and ∼16% inhibition respectively, each greater than their calculated Bliss Additivity, suggesting some synergy. These data demonstrate that non-neutralising mAbs which bind distinct sites in PfRIPR can combine to be growth-inhibitory and better than the expected sum of their parts.

## Discussion

Structural insights into PfPCRCR components and their recognition by parasite growth-inhibitory monoclonal antibodies are already guiding design of improved blood-stage vaccine immunogens. In the case of PfRH5, the most effective antibodies bind around the ‘top half’ of PfRH5, overlapping with or adjacent to the basigin binding site^5,10-12,24^, where they act by sterically blocking binding of PfRH5 to basigin-containing membrane protein complexes^32^. Improved versions of PfRH5 have already been developed^24,33^ and are in clinical trials (NCT05790889 and NCT05978037)^34^, while an immunogen based on the ‘top half’ of PfRH5 is under development. The most growth-inhibitory monoclonal antibodies targeting PfCyRPA also bind to spatially close epitopes, on two blades of the PfCyRPA β-propeller structure^13-15^. Here too, the most effective antibodies act sterically, clashing with the membrane when PfPCRCR approaches basigin on the erythrocyte surface^2^. In contrast to PfRH5 and PfCyRPA, the few known growth-inhibitory antibodies which target PfRIPR bind to sites that are far from the erythrocyte membrane^2,18^, suggesting that they act through a different mechanism. Here we provide the first structural insight into how PfRIPR-targeting antibodies function.

Previous studies mapped the epitopes of all known PfRIPR-targeting growth-inhibitory antibodies to a small region of the tail of PfRIPR containing EGF-like domains 5-8^8,16-18^, but did not reveal how these antibodies work. These EGF-like domains lie distant from both the erythrocyte and merozoite membranes in the centre of the PfRIPR bridge, where our previous structural studies suggested there was flexibility, perhaps between EGF6 and EGF7^2^. We now show that the growth inhibitory antibody RP.012 binds at this location in between EGF6 and 7, where it causes, or stabilises, a substantial bend in the PfRIPR tail. We also show that not all antibodies that bind EGF5-8 are growth-inhibitory and, in contrast to RP.012, non-inhibitory antibodies such as MAD8-2 and MAD8-739 do not stabilise a bent conformation of the PfRIPR tail. This is consistent with a model in which flexibility in the hinge between EGF6 and 7 is required for PfRIPR function (Figure 4C). This flexibility may be required before PfPCRCR binds to basigin on the red blood cell surface, with RP.012 stabilising a conformation of PfPCRCR which is unable to bind basigin. Alternatively, this flexibility might be required to allow the PfPCRCR bridge to bend after basigin binding, allowing the merozoite and erythrocyte membranes to come closer together. RP.012 may prevent the adoption of this specific conformation. Our structural studies suggest that the tail of PfRIPR has multiple such hinge points, including between EGF6 and 7, and between EGF7-8 or EGF8-9, each of which might be targets of growth-inhibitory antibodies.

These findings have consequences for PfRIPR-based immunogen design. The discovery that two of our three EGF5-8 targeting antibodies are not growth-inhibitory cautions against simply including this region of PfRIPR in a vaccine. Indeed, we also reveal a complicated interplay between growth-inhibitory and non-inhibitory antibodies which bind to EGF5-8, showing weak synergy or strong antagonism between antibody pairs. The fact that a non-inhibitory EGF5-8 binding antibody (e.g. MAD8-2) can antagonise the function of a growth-inhibitory antibody (e.g. RP.012), by sterically competing for binding, has consequences for vaccination, as polyclonal sera raised by immunisation with EGF5-8 will likely contain non-inhibitory antibodies which can negatively affect the function of those with growth-inhibition activity. Additionally, these non-inhibitory antibodies may be more frequently raised than growth-inhibitory ones, which need to capture a specific bent conformation of EGF-like domains. A better immunogen might therefore include EGF6-7 locked in the bent form stabilised by RP.012 to specifically induce more growth-inhibitory responses.

We also strikingly demonstrate that antibodies which bind to spatially distinct sites of PfRIPR can act synergistically, as MAD8-66, which binds to PfRIPR C-terminal domain, synergises with EGF5-8-binding antibodies. This is observed most strongly for MAD8-66 and growth-inhibitory RP.012. It is also seen, albeit more weakly, for MAD8-2 and MAD8-739. The mechanism by which synergy occurs is currently not clear. However, it seems likely that whatever conformational changes take place in PfRIPR during erythrocyte invasion are incompatible with the presence of antibodies bound simultaneously to the CTD and to EGF5-8. This is perhaps due to steric effects with direct clashes of these antibodies with one another, or with other parts of PfRIPR. Multiple antibodies might also be able to interfere with the conformation of PfRIPR tail and its many hinge points more effectively than single antibodies. Alternatively, MAD8-66 might disrupt the interface of PfRIPR and PfPTRAMP, changing how PfPCRCR is presented on the parasite surface, despite not leading to its disassembly. Future studies will reveal these mechanisms, using larger panels of human antibodies.

Our study therefore has multiple consequences for PfRIPR-based vaccine immunogen design. A vaccine candidate comprising a fusion of PfCyRPA and EGF-like domains 7-8 of PfRIPR is currently in clinical trials in combination with the blood-stage vaccine RH5.1 (NCT05385471). This immunogen lacks both the epitope for growth-inhibitory antibody RP.012 and the epitope for the synergistic antibody MAD8-66. Rational design of future PfRIPR-based vaccine immunogens is essential, and the best immunogen may require discontinuous regions of PfRIPR, including both EGF5-8 and the CTD. The discovery that growth-inhibitory and antagonistic antibodies bind overlapping epitopes on EGF5-8 presents a further challenge for immunogen design, suggesting that protein engineering will be required to generate immunogens which specifically elicit only growth-inhibitory antibodies and those which synergise with them, without raising antagonistic antibodies.

Future studies of large panels of human growth-inhibitory antibodies targeting PfRIPR will be important to provide deeper insight into the nature of these growth-inhibitory epitopes and the conformation of PfRIPR when bound to these antibodies. This will allow rational design of the most effective PfRIPR-based vaccine immunogen for next-generation malaria vaccines.

## Limitations of the study

A limitation of this study is the relatively small size of our antibody panel, containing just four antibodies. This panel also does not contain a human growth-inhibitory antibody, but we do not expect that growth inhibitory antibodies such as RP.012 are unique to mice. Given the large molecular weight and the elongated structure of PfRIPR, the epitopes of our antibodies cover only a small amount of its total available surface. Larger panels of human antibodies will be required to obtain a more complete understanding of the human antibody response to PfRIPR and the growth inhibition potential of its entire surface. However, this will require a substantial effort, and the findings of this study are already informative and timely. Our findings demonstrate that critical epitopes of PfRIPR are not currently included in a PfRIPR-based vaccine in clinical trial, and these findings also provide enough information to guide rational design of new PfRIPR immunogens. Future larger antibody panels will continue to inform this immunogen design.

## Resource availability

### Lead contact

Requests for further information and resources should be directed to and will be fulfilled by the lead contact MKH.

### Materials availability

Unique reagents generated in this study will be made available on request by the lead contact with a completed Materials Transfer Agreement.

### Data and code availability

Crystallographic data and protein models are available from the Protein Data Bank (PDB) under accession codes 9TUW for RP.012-bound PfRIPR, 9TUX for MAD8-66-bound PfRIPR, and 9TUY for MAD8-739-bound PfRIPR. Cryo-EM maps and associated co-ordinates for MAD8.2-bound PfRIPR are available from the Electron Microscopy Data Bank (EMDB) under accession code 9TUZ and EMDB-56281, and for PfRIPR-PfCyRPA at 9TV0 and EMDB-56282. A previously published structure and cryo-EM map have been used in this study; these can be found in the PDB under accession codes 8CDD and the EMDB at EMDB-16640. All other data is available from the authors on request.

## Acknowledgements

This work was funded through a Wellcome Investigator award (220797/Z/20/Z), a Medical Research Council Award (MR/Z505687/1) and a Gates Foundation Award (INV-078815). We thank Barnabas G. Williams and Simon J. Draper (University of Oxford) for providing expression constructs for RP.012 and PfRIPR EGF5-8. We thank Boubacar Traore (University of Sciences, Technique and Technology of Bamako, Mali) and Peter Crompton (NIAID) for providing PBMC samples, Simon J. Draper (University of Oxford) for providing recombinant PfRIPR, Gavin Wright (University of York) for providing biotinylated CD4 and Ludmila Krymskaya for sorting PBMC samples. We thank Rishi Matadeen and Edward Lowe at the COSMIC facility (University of Oxford) for support with cryo-EM data collection and data processing, and Edward Lowe for support with crystallography. We thank beamline scientists at Diamond Light Source beamlines I04 and I24. We thank Rod Chalk (University of Oxford) for support with mass spectrometry and David Staunton (University of Oxford) for support with surface plasmon resonance. We also thank Carole Long and Ababacar Diouf for support with growth inhibition assays. We thank Jacqueline Kirchner, Holger Kanzler, Hedda Wardemann and Annie Zumsteg from the Gates Foundation for interesting discussion. We thank the NHS Blood and Transplant Services for the supply of human blood for parasite culture and growth inhibition assays. This research was supported in part by the Intramural Research Program of the National Institutes of Health (NIH). The contributions of the NIH author(s) are considered Works of the United States Government. The findings and conclusions presented in this paper are those of the author(s) and do not necessarily reflect the views of the NIH or the U.S. Department of Health and Human Services.

## Author contributions

B.F. expressed and purified proteins, performed SPR analysis, FRET analysis, and X-ray crystallographic structure determination. A.J.R.C., L.T.W. and J.T. isolated and characterised human antibodies. B.F. and E.E-W. conducted cryo-EM structure determination and antibody competition analysis. E.B. performed GIA assays. J.T. and M.K.H. designed experiments and contributed expertise and funding. B.F., A.J.R.C., J.T. and M.K.H. prepared the manuscript and all authors contributed and commented.

## Supplementary information titles and legends

Figures S1-5, Tables S1-7

## STAR Methods

### Key resources table

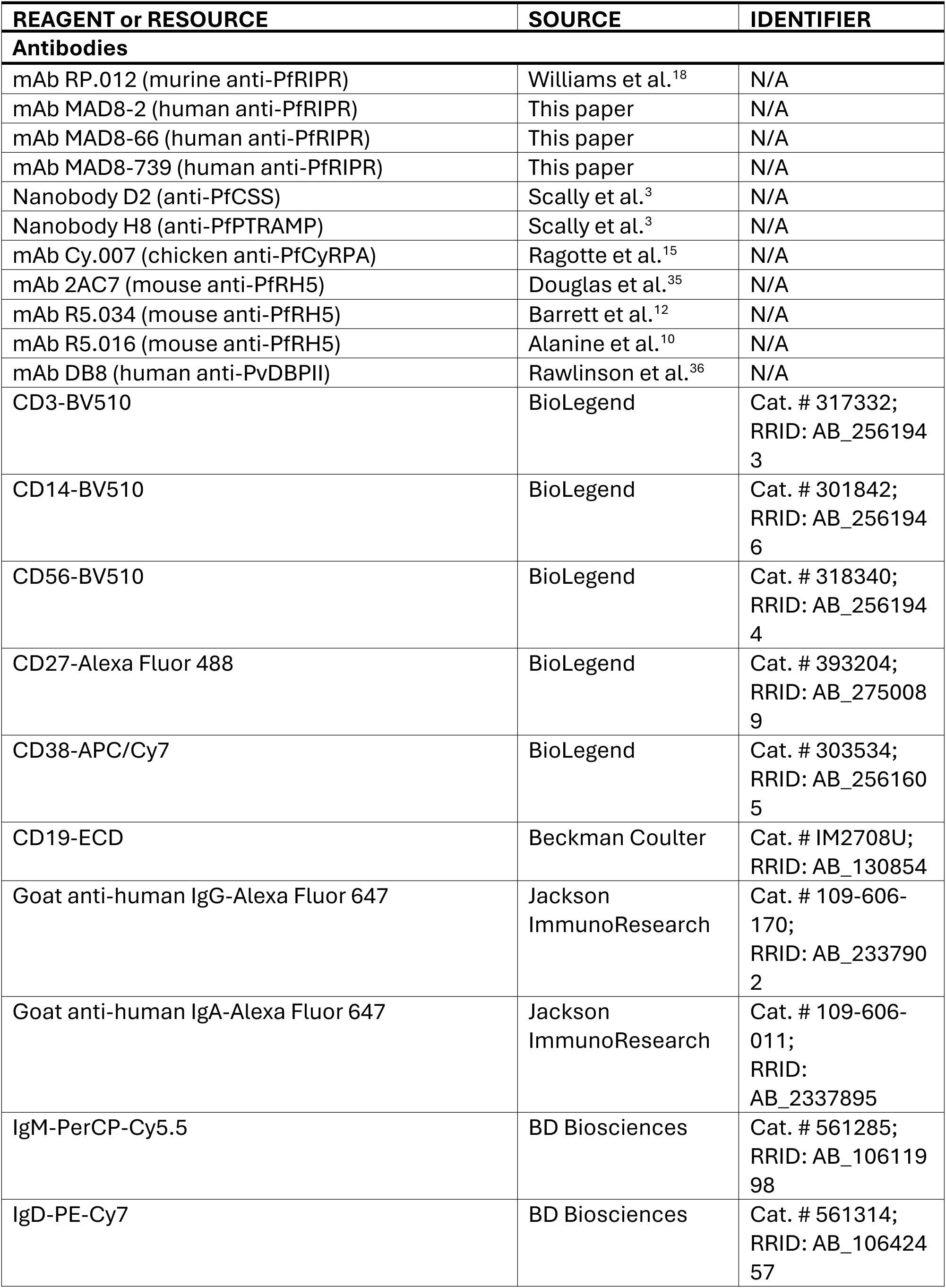
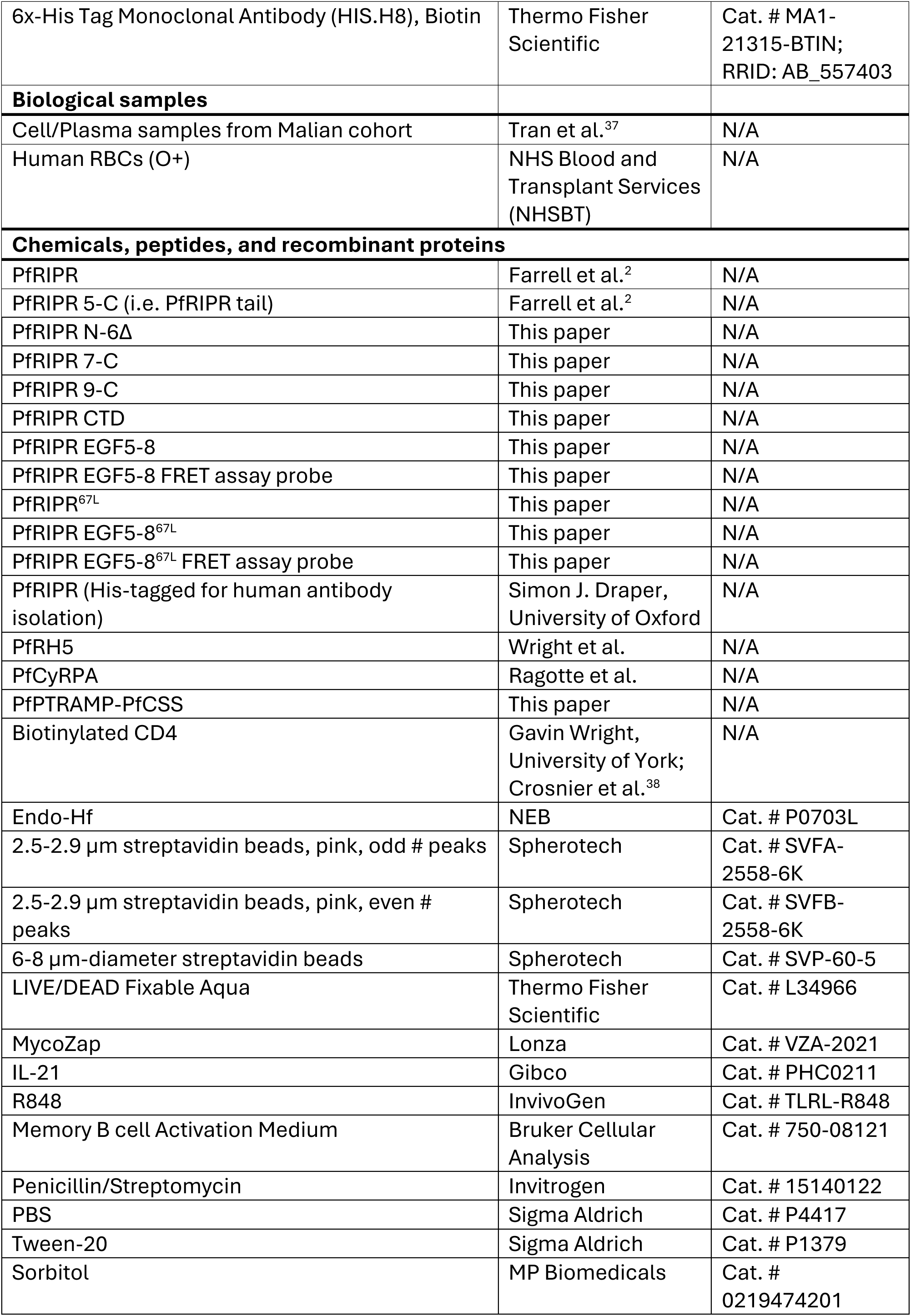
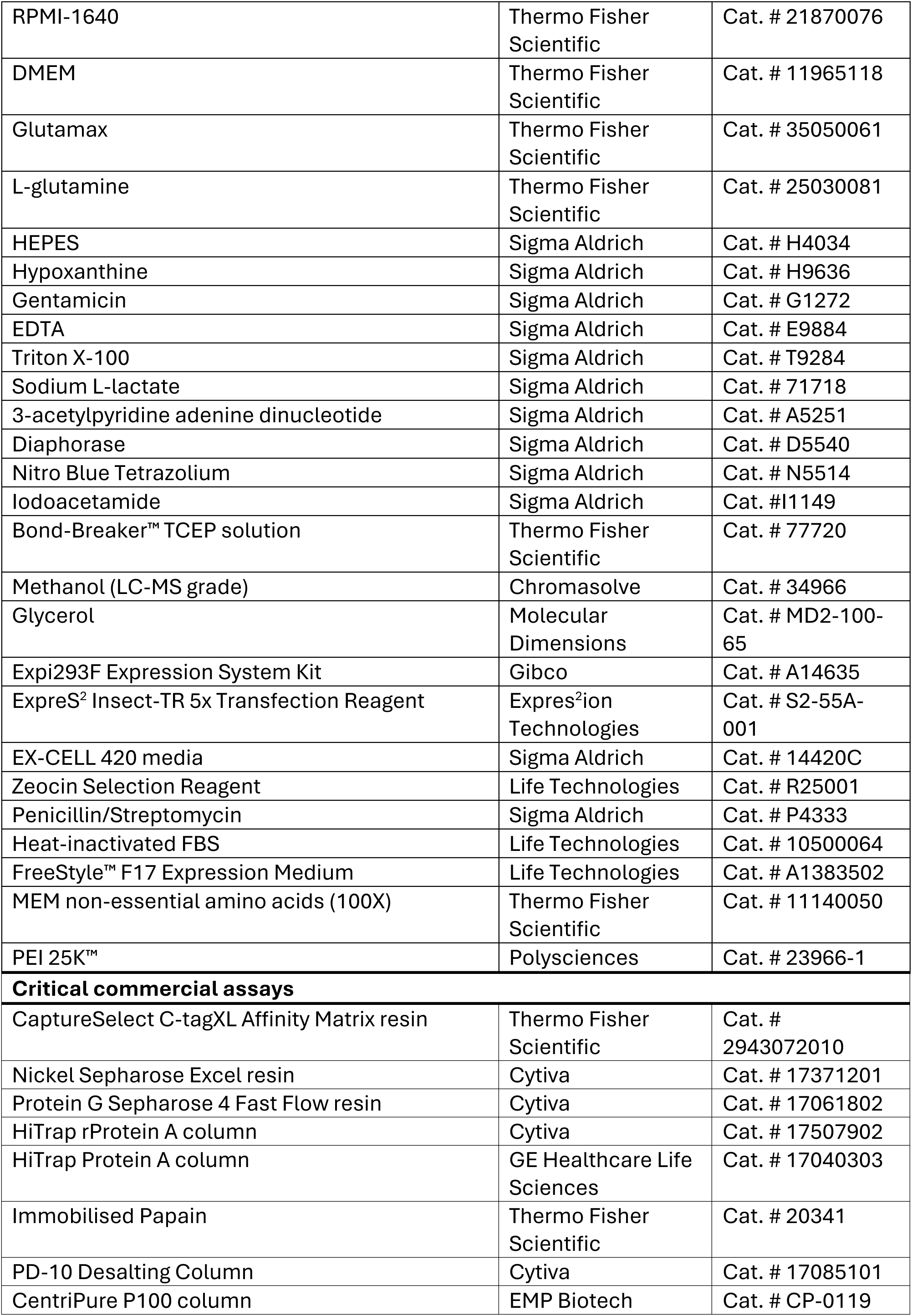
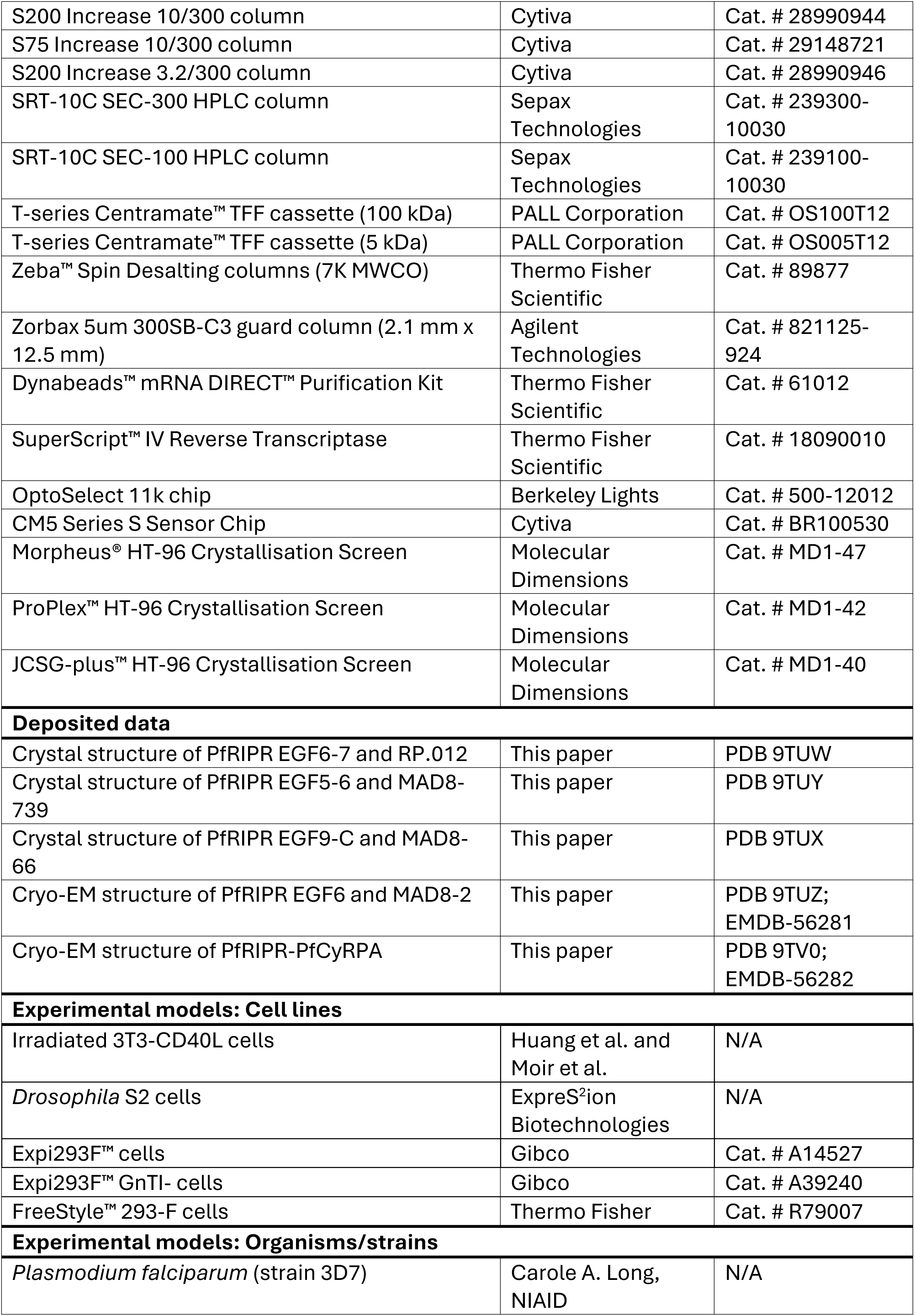
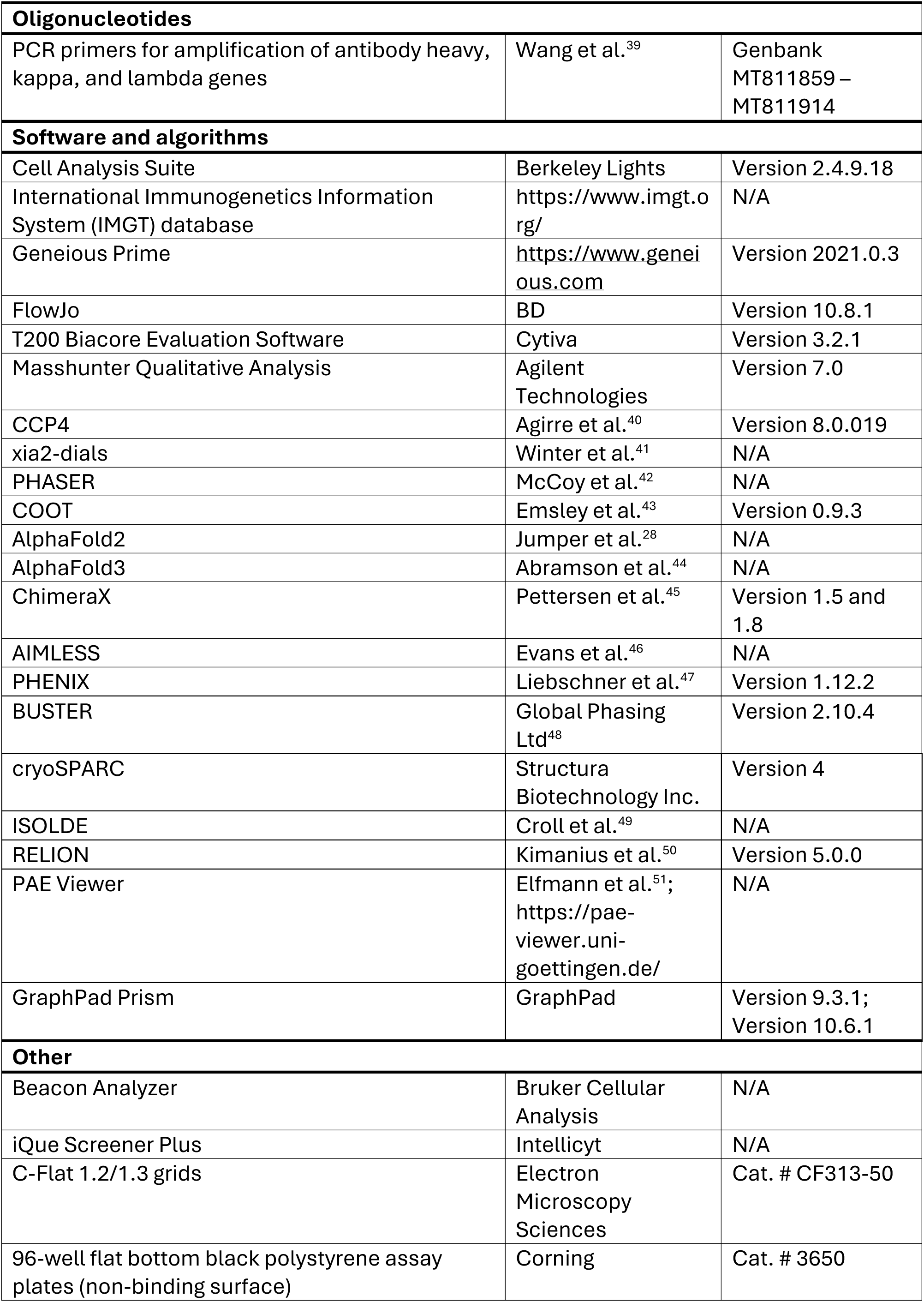

### Experimental model and study participant details

#### Human studies

Peripheral blood mononuclear cells (PBMCs) were sourced from a cohort of 758 malaria-exposed donors from ages 1 month to 41 years in the village of Kalifabougou, Mali^37,52^. Samples were taken from subjects several times a year before and during the malaria season. The Kalifabougou cohort study is an ongoing longitudinal study approved by the Ethics Committee of the Faculty of Medicine, Pharmacy and Dentistry at the University of Sciences, Technique and Technology of Bamako, and the Institutional Review Board of the National Institute of Allergy and Infectious Diseases, National Institutes of Health (NIH IRB protocol number: 11IN126; https://clinicaltrials.gov/; trial number NCT01322581). Written informed consent was obtained from all participants, or from parents or guardians of child participants, before inclusion in the study.

#### Cell lines

Expi293F™ cells (Gibco) and Expi293F™ GnTI- cells (Gibco) were cultured in Expi293F™ Expression Medium at 37°C, 8% CO_2_ and 70% humidity, shaking at 110 rpm. Each were transfected using the Expi293F Expression System (Gibco, Thermo Fisher Scientific) according to the manufacturer’s instructions. FreeStyle™ 293-F cells (Thermo Fisher) were cultured in FreeStyle™ F17 Expression Medium containing L-glutamine and 1 x MEM non-essential amino acids (Gibco) at 37°C, 8% CO_2_ and 70% humidity, shaking at 110 rpm. Cells were transfected using a 3:1 ratio of PEI (PEI 25K™, Polysciences) and DNA. *Drosophila* S2 cells (ExpreS^2^ions Biotechnologies) were cultured in EX-CELL 420 medium (Sigma Aldrich) containing 100 U/mL penicillin and 0.1 mg/mL streptomycin at 25°C, shaking at 120 rpm. S2 cells were transfected as static cultures at 25°C using ExpreS^2^ Insect-TR 5x Transfection Reagent (ExpreS^2^ion Biotechnologies) according to the manufacturer’s instructions, and stable polyclonal cell lines were selected using Zeocin Select Reagent (Life Technologies). For generation of feeder cells, 3T3-CD40L cells were cultured in Glutamax-supplemented DMEM (Thermo Fisher Scientific) with 10% FBS and 100 U/mL penicillin and 0.1 mg/mL streptomycin at 37°C and 5% CO_2_ and irradiated at 5000 rads for 5 minutes and 30 seconds. The human IgG1 antibodies were expressed in the HD293F cell line (Genscript).

### Method Details

#### Production of PfRIPR constructs

PfRIPR was expressed and purified as described previously^2^. PfRIPR (D21-N1086) with a C-terminal C-tag was recombinantly expressed using a polyclonal stable S2 cell line (ExpreS^2^ion Biotechnologies). This PfRIPR construct and all others in this study have N to Q substitutions at residues 103, 144, 228, 303, 334, 480, 498, 506, 526, 646, 647, 964 and 1021 to remove potential N-linked glycosylation sites. Cultures were harvested after 3-4 days and PfRIPR was purified from the supernatant using CaptureSelect C-tagXL Affinity Matrix resin (Thermo Fisher) as recommended, after concentration and buffer exchange into 50 mM Tris pH 7.5, 150 mM NaCl by tangential flow filtration with 100 kDa cassettes. Eluted proteins were desalted into 20 mM HEPES pH 7.5, 150 mM NaCl (HBS) using a PD-10 desalting column (Cytiva). PfRIPR tail (i.e. PfRIPR 5-C, D717-N1086) was expressed and purified as for PfRIPR except that tangential flow filtration was performed with 5 kDa cassettes. Further PfRIPR constructs (PfRIPR N-6Δ (D21-N816 except T482-I555), PfRIPR 7-C (D817-N1086), PfRIPR 9-C (E899-N1086) and PfRIPR CTD (N981-N1086) were expressed and purified as PfRIPR tail. Following affinity purification, proteins were further purified by gel filtration using either a S200 Increase 10/300 or S75 Increase 10/300 column as appropriate into HBS pH 7.5.

PfRIPR EGF5-8 (D717-D900) with an N-terminal monoFc tag and TEV protease cleavage site and a C-terminal C-tag was recombinantly expressed in Expi293F™ cells (Gibco) as recommended. Culture supernatants were harvested then filtered using 0.45 µm filters before being applied to Protein G Sepharose 4 Fast Flow resin (Cytiva). Following washing with 20-30 column volumes of HBS, bound proteins were eluted with 0.1 M glycine pH 2.0, immediately neutralised with Tris pH 8.0, then buffer exchanged into HBS. To remove the monoFc tag, buffer-exchanged proteins were incubated with TEV protease overnight at room temperature, then re-applied to Protein G resin. PfRIPR EGF5-8 from the unbound fraction was collected, then bound proteins were recovered by elution as before. PfRIPR EGF5-8 was further purified by gel filtration using a S200 Increase 10/300 column (Cytiva) equilibrated in HBS pH 7.5.

A FRET probe^29^ comprising mCerulean-(GGS)_2_-PfRIPR(EGF5-8)-(GGS)_2_-mCitrine with a C-terminal His-tag was expressed in Expi293F™ cells as recommended. Culture supernatants were harvested then filtered using 0.45 µm filters before being applied to Nickel Sepharose Excel resin (Cytiva). After washing with 20-30 column volumes of 25 mM Tris pH 8.0, 150 mM NaCl (TBS) and then 5 column volumes of 25 mM Tris pH 8.0, 500 mM NaCl, 20 mM imidazole, bound proteins were eluted using 5 column volumes of 25 mM Tris pH 8.0, 150 mM NaCl, 500 mM imidazole. The PfRIPR FRET probe was further purified by gel filtration using a S200 Increase 10/300 column (Cytiva) equilibrated in HBS pH 7.5 as above.

A variant of PfRIPR containing a designed disulphide bond between EGF-like domains 6 and 7 was generated by introducing the substitutions H804C and F835C. Versions of full-length PfRIPR (PfRIPR^67L^), PfRIPR EGF5-8 (PfRIPR EGF5-8^67L^) and the PfRIPR FRET probe were expressed and purified as for their wild-type versions.

#### Production of PfPCRCR components

PfRH5 (K140-Q526, i.e. PfRH5ΔN^5^) was expressed and purified as for PfRIPR tail and as detailed previously^2^. This PfRH5 construct additionally contains the substitutions C203Y (for the 7G8 *P. falciparum* strain) and T216A and T299A to remove potential N-linked glycosylation sites, and a C-tag. PfCyRPA (D29-E362 with substitutions S147A, T324A and T340A, and a C-tag) was expressed in Expi293F™ cells and purified using CaptureSelect C-tagXL Affinity Matrix resin (Thermo Scientific) as detailed previously^2,15^. PfPTRAMP-PfCSS was recombinantly expressed by co-transfecting Expi293F™ GnTI- cells (Gibco) for co-expression of PfPTRAMP (L42-T307 with a C-tag) and PfCSS (Q21-K290 with a C-terminal Avi-His-tag). PfPTRAMP-PfCSS heterodimer was purified by tandem affinity purification using Nickel Sepharose Excel resin (Cytiva), then following desalting into HBS pH 7.5, by additional purification with CaptureSelect C-tagXL Affinity Matrix resin (Thermo Scientific). Purified PfPTRAMP-PfCSS heterodimer was deglycosylated by incubation with Endo-Hf (NEB) overnight at 37°C, then further purified by gel filtration using a S200 Increase 10/300 into HBS pH 7.5.

#### Isolation of human antibodies against PfRIPR

Recombinant His-tagged PfRIPR (provided by Prof Simon Draper, University of Oxford) was conjugated to 2.5-2.9 µm-diameter streptavidin beads (Spherotech) via a biotinylated anti-His-tag antibody (Thermo Fisher Scientific). Beads were pre-incubated with this capture antibody for 20 minutes at room temperature before incubation with 10 µg/mL PfRIPR for 1 hour, also at room temperature. Washes were carried out with 0.5% BSA-PBS, and conjugated beads were blocked with 10 µg/mL biotinylated CD4 (provided by Gavin Wright, University of York^38^) to saturate streptavidin sites on beads.

PfRIPR-specific B cells were isolated using an optofluidics method as previously described^11^. Briefly, cryopreserved PBMCs from Kalifabougou were thawed and stained in PBS at 4°C for 20 minutes with LIVE/DEAD Fixable Aqua (Thermo Fisher Scientific) to specifically stain dead cells. Cells were subsequently stained in 1% FBS-PBS at 4°C for 20 minutes with the following panel of fluorescent antibodies: CD3-BV510, CD14-BV510, CD56-BV510, CD27-Alexa Fluor 488, CD38-APC/Cy7 (BioLegend), CD19-ECD (Beckman Coulter), IgA-Alexa Fluor 647 (Jackson ImmunoResearch), IgM-PerCP-Cy5.5, and IgD-PeCy7 (BD Biosciences). IgG^+^ memory B cells in stained PBMC samples were classified as live CD3^−^ CD14^−^ CD56^−^ CD19^+^ IgD^−^ IgM^−^ IgA^−^ singlets. This cell population was sorted and cells were cultured according to one of two methods. In method A, cells were seeded at 2,500/mL in U-bottom 96-well plates in Memory B cell Activation Medium (Bruker Cellular Analysis) for 6 days. Method B involved seeding cells at 10,000-25,000/mL in 384-well plates in I10 media (Iscove’s modified Dulbecco’s Medium, 10% FBS, 1:1000 MycoZap; Thermo Fisher Scientific and Lonza) supplemented with 100 ng/mL IL-21 (Gibco) and 0.5 mg/mL R848 (InvivoGen), with 300,000/mL irradiated CD40L^+^ 3T3 feeder cells^53,54^, for 10 days. One day before culture conclusion, culture supernatants were sampled and screened using PfRIPR-conjugated streptavidin beads previously described and polyclonal anti-IgG secondary antibody (Jackson ImmunoResearch). On the last day of culture (Day 6 for culture method A or Day 10 for culture method B), B cells in positive wells identified the previous day were resuspended in fresh I10 media, again supplemented with IL-21 and R848, and loaded into the channels of an OptoSelect 11k chip. These cells were then directed into nanolitre-volume pens using opto-electropositioning (OEP) light cages in a Beacon Cell Analysis Suite (Bruker Cellular Analysis). Unpenned cells were flushed from channels. Channels were then flooded with 6-8 µm-diameter streptavidin beads (Spherotech) pre-conjugated to 1 µg/mL PfRIPR. These beads were imported with polyclonal anti-IgG secondary antibody (Jackson Immunoresearch). B cells with PfRIPR-specific antibody production were then identified by the appearance of fluorescent ‘blooms’ in channels. These cells were exported to chilled lysis buffer and frozen at -80°C until PCR amplification and sequencing was performed.

#### mAb sequence analysis and production

PCR amplification and sequencing of antibodies was performed as previously described^11,39^. Briefly, B cell lysates were thawed and subjected to RNA cleanup using Dynabeads™ mRNA DIRECT™ Purification Kit (Thermo Fisher), and then RNA was reverse transcribed to cDNA using SuperScript™ IV Reverse Transcriptase (Thermo Fisher). cDNA for heavy and light chain variable regions was specifically amplified using a cocktail of primers (primer sequences available in GenBank MT811859 to MT811914). Sequence analysis of heavy and light chain variable regions was performed with reference to the International Immunogenetics Information System (IMGT) database, yielding sequence annotations, predicted genetic background, and antibody isotype^55^. Sequences were cloned into plasmids containing an IgG1 or relevant light chain backbone (GenScript) and transfected into Expi293F™ cells (Thermo Fisher Scientific). HiTrap Protein A columns (GE Healthcare Life Sciences) were used to purify recombinant IgG. PfRIPR-binding activity of the recombinant mAbs was confirmed using antigen-coated streptavidin beads.

#### Expression of antibodies and nanobodies

Monoclonal antibodies (RP.012, MAD8-2, MAD8-66 and MAD8-739) were expressed in Expi293F™ cells as recommended by co-transfection of heavy and light chain plasmids. The murine antibody RP.012 was recombinantly expressed as a chimeric human antibody composed of the mouse variable domain and human IgG1 constant domains. Human antibodies MAD8-2, MAD8-66 and MAD8-739 were expressed as human IgG1. Culture supernatants were harvested, then filtered with 0.45 µm filters before applying to Protein G Sepharose 4 Fast Flow resin (Cytiva). Following washing with 20-30 column volumes of phosphate buffered saline (PBS, #P4417 Sigma Aldrich), antibodies were eluted using 0.1 M glycine pH 2.0, then immediately neutralised with Tris pH 8.0 and buffer exchanged into PBS using Amicon Ultra centrifugal filters.

Fab fragments of RP.012, MAD8-2, MAD8-66, MAD8-739 were recombinantly expressed in Expi293F™ cells. In this case, RP.012 was expressed with its mouse constant domain. All constructs had a C-terminal His-tag on the heavy chain and were purified as for the PfRIPR FRET probe. Fab fragments of the PfCyRPA Cy.007 antibody^15^ were obtained by expressing the monoclonal antibody as above followed by cleavage with immobilised papain (#20341, Thermo Scientific) and separation of the Fab fragments by using a 1 mL HiTrap rProtein A prepacked column (Cytiva).

Single chain variable fragments (scFv) of RP.012 and MAD8-739 were expressed using Expi293F™ cells. These were constructed by joining the variable heavy (VH) and variable light (VL) domains of each antibody with a (GGGGS)_3_ linker in the format VH-(GGGGS)_3_-VL. Each scFv fragment was purified as for Fab fragments, then further purified by gel filtration using a S200 Increase 10/300 column (Cytiva) in HBS pH 7.5 to isolate the monomeric fraction.

The anti-PfPTRAMP nanobody H8^3^ and the anti-PfCSS nanobody D2^3^ were expressed in FreeStyle™ 293-F cells (Thermo Fisher) in FreeStyle™ F17 Expression Medium containing L-glutamine and 1 x MEM non-essential amino acids (Gibco). Cells were transfected using a 3:1 ratio of PEI (PEI 25K™, Polysciences) and DNA. After 6 days, culture supernatants were harvested and 0.45 µm filtered, then their pH adjusted to pH 8.0 using Tris before applying to Nickel Sepharose Excel resin (Cytiva). Following washing with 30 column volumes of TBS pH 8.0, bound nanobodies were eluted with 25 mM Tris pH 8.0, 150 mM NaCl, 500 mM imidazole. Eluted proteins were exchanged into PBS using a CentiPure P100 column (EMP Biotech), then further purified by gel filtration using an SRT-10C SEC-100 HPLC column (Sepax Technologies) in PBS.

#### Epitope binning of mAbs by size exclusion chromatography

Purified Fab fragments of each antibody (RP.012, MAD8-2, MAD8-66 or MAD8-739) were mixed with various fragments of PfRIPR at equimolar concentration and incubated for 5 minutes at room temperature. Binding analysis was performed by injecting each mixture on a S200 Increase 10/300 or a S200 Increase 3.2/300 column (Cytiva) equilibrated in HBS pH 7.5. Peak fractions were analysed by SDS-PAGE to determine whether proteins co-eluted. Additionally, PfRIPR tail, PfRIPR 9-C and Fab fragments were injected alone to verify co-elution where necessary.

#### Assessment of competition between mAbs and PfPCRCR components

To assess whether antibodies could bind to the PfRCR complex, PfRCR was reconstituted by mixing PfRH5 and PfCyRPA at 1.1x molar excess over PfRIPR for 5 mins at room temperature. This was concentrated using a 30K MWCO Amicon Ultra centrifugal unit, then injected onto a S200 Increase 10/300 column (Cytiva) equilibrated in HBS pH 7.5. Fractions corresponding to the PfRCR complex were combined and concentrated using a 30K MWCO Amicon Ultra centrifugal unit at 10,000 g and 4°C to 0.8 mg/mL. 5 µg of Fab fragments for each antibody (RP.012, MAD8-2, MAD8-66 and MAD8-739) were incubated with 10 µg purified PfRCR complex (equating to ∼4-fold molar excess of Fab fragment) for 5 minutes at room temperature. Samples were then injected on a S200 Increase 3.2/300 column (Cytiva) in HBS pH 7.5 and peak fractions analysed by SDS-PAGE. 5 µg of Fab fragments and 10 µg of PfRCR complex alone were injected to verify peak shifting and co-elution of all components.

To assess whether antibodies could bind the PfPTRAMP-PfCSS-PfRIPR (PfPCR) complex, mixes were prepared containing 2 µM PfRIPR tail (4.5 µg), 2 µM Fab fragments (5 µg), and

5.5 µM PfPTRAMP-PfCSS, then incubated at room temperature for 5 minutes. PfPTRAMP-PfCSS was used at this ∼2.75x molar excess such that it was present at a concentration above its binding affinity for PfRIPR^2,3^. Samples were injected on a S200 Increase 3.2/300 column (Cytiva) in 20 mM HEPES pH 7.5, 50 mM NaCl then peaks analysed by SDS-PAGE. This was first performed for MAD8-2. PfPTRAMP-PfCSS, PfRIPR^tail^, PfRIPR^tail^+MAD8-2, and MAD8-2 alone were injected to verify whether components co-elute in a ternary complex. Samples for RP.012, MAD8-66 and MAD8-739 were compared to data for the MAD8.2-bound complex.

#### FRET analysis

The PfRIPR FRET probe (wild-type or EGF6-7 cysteine locked) was diluted to 50 µL at 1 µM using HBS pH 7.5 in a 96-well flat bottomed black polystyrene assay plate (non-binding surface, #3650, Corning) and the base level emission spectrum was recorded over 455-625 nm (with a 10 nm bandwidth) after excitation at 420 nm (with a 10 nm bandwidth) using a CLARIOstar plate reader (BMG LabTech). The gain was adjusted to 50% using emission at 475 nm. Fab fragments or an equivalent volume of buffer were added to a final 3 µM concentration and the emission spectrum recorded once more. FRET efficiency was quantified by calculating the FRET ratio by dividing the fluorescence emission intensity at 525 nm by that at 475 nm.

#### Surface Plasmon Resonance analysis

To measure binding kinetics of antibodies, purified mAbs (chimeric RP.012, MAD8-2, MAD8-66 and MAD8-739) were prepared at 10 nM in SPR buffer (20 mM HEPES pH 7.5, 150 mM NaCl, 0.01% Tween 20) then captured by Protein A/G immobilised on a CM5 Series S Sensor chip for 30 seconds at 5 µL/min. PfRIPR (either wild-type or RIPR^67L^) was diluted in SPR buffer to 100 nM and a two-fold dilution series over six steps was prepared in the same buffer. Binding was assessed by recording SPR traces on a T200 Biacore Instrument (Cytiva) in SPR buffer at 25°C, injecting each sample for 120 seconds at 30 µL/min and allowing dissociation for a further 240 seconds. Between sample injections, the chip was regenerated by injecting 100 mM glycine pH 2.0 for 60 seconds at 10 µL/min, then the appropriate mAb was recaptured as before. Experiments were recorded twice each on two different days, equalling a total of 4 independent measurements. Kinetic parameters were estimated using T200 Biacore Evaluation Software (Cytiva) by fitting data to a global 1:1 binding model. The binding affinity and kinetic parameters are expressed as the mean of these fitted parameters. One SPR trace each for 100 nM PfRIPR against MAD8-66 and MAD8-739 was excluded from analysis due to abnormal antibody capture in these cycles. The interaction between PfRIPR^67L^ and RP.012 chimeric mAb was measured only three times.

To conduct competition binning of antibodies, Fab fragments (RP.012, MAD8-2, and MAD8-739) were prepared at 1 µM in SPR buffer (20 mM HEPES pH 7.5, 150 mM NaCl, 0.01% Tween 20) then injected sequentially over a CM5 Series S Sensor chip (Cytiva) with PfRIPR immobilised on its surface by standard amine coupling as detailed previously^2^. SPR traces were recorded as above except using a flow rate of 5 µL/min, injecting the first Fab fragment for 120 seconds, waiting 30 seconds, then injecting the second Fab fragment for 120 seconds. Dissociation was then allowed for 120 seconds. After each injection cycle of two sequential Fab fragments, the chip was regenerated using 100 mM glycine pH 2.0 or 50 mM NaOH where appropriate. Each competition injection cycle was measured in triplicate using Fab fragments prepared independently.

#### Growth inhibition activity assay for antibodies and fragments

The 3D7 strain of *P. falciparum* was cultured in O+ RBCs obtained from NHS Blood and Transplant services (NHSBT). On day 1 of the assay, 30 mL of culture were synchronised twice with 5% sorbitol (MP Biomedicals) and cultured at 37°C overnight in complete medium (RPMI (Life Technologies), 1% L-glutamine (Life Technologies), 25 mM HEPES (Sigma), 0.005% hypoxanthine (Sigma), 10% heat-inactivated pooled O+ human serum (NHSBT), 10 µg/mL gentamicin (Sigma)) at 2% haematocrit. The following morning, late-stage parasites were diluted to 0.4% parasitaemia in 2x complete medium (RPMI, 1% L-glutamine, 25 mM HEPES, 0.005% hypoxanthine, 20% heat-inactivated pooled O+ human serum, 20 µg/ml gentamicin) at 2% haematocrit. Twenty µl of the parasitised RBC suspension were added to each well of 96-well half area plates containing 20 µl of 10 mM EDTA (100% inhibition control; Sigma), incomplete medium (0% inhibition control; RPMI, 1% L-glutamine, 25 mM HEPES, 0.005% hypoxanthine), monoclonal antibody controls provided to give ∼50% inhibition (2AC7^35^ (40 µg/ml), R5.034^12^ (5 µg/ml), R5.016^10^ (30 µg/ml)) monoclonal antibody negative control (DB8^36^ (1000 µg/ml)) or monoclonal antibodies, Fab or scFv fragments of interest, or their combinations in incomplete medium. The plates were then transferred to a modular incubator chamber, gassed with 5% O2, 5% CO2, 90% nitrogen and incubated at 37°C for 44 h. A parallel tracker culture at 0.4% parasitaemia was used to monitor parasite growth.

On day 4 of the assay, the plates were removed from the incubator and washed twice with 100 µL cold PBS per well. Parasitised RBCs were resuspended by shaking for 1 min at maximum speed on Titramax 100 (Heidolph). Parasite growth was estimated using the lactate dehydrogenase (LDH) assay: 120 µl of LDH substrate (containing 100 mM Tris, pH 8.0, 0.25% Triton X-100 (Sigma), 50 mM sodium L-lactate (Sigma), 3-acetylpyridine adenine dinucleotide (50 μg/ml; Sigma), diaphorase (1 U/ml; Sigma) and nitro blue tetrazolium (0.2 mg/ml; Sigma)) was added per well and optical density (OD) readings were taken at 650 nm when OD rates in 0% inhibition control wells reached 0.4-0.6.

Percent inhibition was calculated using the following formula:

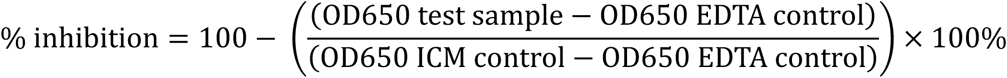

Monoclonal antibodies were assayed at final 1 mg/mL concentration or at combinations of 1 mg/mL each. The Fab fragment and scFv fragment of RP.012 were assayed at twice the molar concentration of the monoclonal of RP.012 to match the molar amounts of paratope. All GIA assays were performed on the same plate allowing direct comparison between antibodies, their fragments and combinations. To identify synergy and antagonism, the predicted Bliss Additivity^31^ growth inhibition values for each antibody combination was calculated as reported previously^56^.

#### Mass Spectrometry of EGF6-7 cysteine-locked PfRIPR

To confirm the formation of disulphide bonds in the locked EGF6-7 protein, PfRIPR EGF5-8 (wild-type and cysteine-locked between EGF6 and 7) were exchanged into and diluted to 0.5 mg/ml with denaturing buffer (200 mM ammonium bicarbonate pH 8, 6 M guanidine hydrochloride) using Zeba™ Spin Desalting columns (7K MWCO, Thermo Fisher). 10 µg of each sample was incubated at 50°C for 30 minutes, then the denatured samples cooled on ice. To each, freshly prepared iodoacetamide was added to a final 20 mM concentration and cysteine alkylation allowed to proceed for 30 minutes at room temperature in the dark. Samples were then exchanged into native buffer (200 mM ammonium bicarbonate pH 8). For samples to be labelled with iodoacetamide under reducing conditions, samples were supplemented with 5 mM TCEP (Bond-Breaker™ TCEP Solution, Thermo Fisher) prior to protein denaturation at 50°C. To act as an unreacted control, separate samples were also exchanged directly into native buffer.

Intact mass spectrometry was used to detect the degree of iodoacetamide-induced cysteine alkylation of each of these PfRIPR EGF5-8 samples under reducing and non-reducing conditions, given that each alkylation event introduces a carbamidomethyl group of 52.07 Da. Reversed-phase chromatography was performed in-line prior to mass spectrometry using an Agilent 1290 uHPLC system (Agilent Technologies Inc.). Protein samples were diluted to 0.02 mg/ml in 0.1% formic acid and 50 µl was injected on to a 2.1 mm x 12.5 mm Zorbax 5um 300SB-C3 guard column housed in a column oven set at 40°C. Proteins were eluted using a solvent system consisting of 0.1% formic acid in ultra-high purity water (solvent A) and 0.1 % formic acid in methanol (LC-MS grade, Chromasolve) (solvent B), first using a linear gradient of 10% to 80% solvent B over 35 seconds at a flow rate of 1.0 ml/min, then isocratically at 95% solvent B for 40 seconds. This was then followed by equilibration at the initial solvent condition for 15 seconds. Protein intact mass was determined using a 6530 electrospray ionisation quadrupole time-of-flight mass spectrometer (Agilent Technologies), configured with the standard electrospray ionisation source and operated in positive ion mode. The ion source was operated with a capillary voltage of 4000 V, nebuliser pressure of 60 psig, and drying gas at 35°C, flowing at 12 L/min. The instrument ion optic voltages were: fragmentor 250 V, skimmer 60V, and octupole RF 250 V. Data analysis was performed using the Masshunter Qualitative Analysis v7.0 (Agilent Technologies).

No free cysteines were detected in either wild-type EGF5-8 or EGF5-8^67L^ in non-reducing conditions, with observed masses matching their calculated masses and the unreacted controls. By contrast, 26 free cysteines were detected for wild-type EGF5-8 and 28 for EGF5-8^67L^ under reducing conditions. Together, these data are consistent with EGF5-8^67L^ containing 13 native and one additional disulphide bond as designed, validating successful introduction of a cysteine lock between EGF6 and 7.

#### Structure determination by X-ray crystallography

A complex of the Fab fragment of RP.012 and PfRIPR EGF6-7 was isolated by gel filtration using a S75 Increase 10/300 column in HBS pH 7.5. Crystals were obtained using sitting drop vapour diffusion by mixing 100 nL reservoir solution with 100 nL protein complex at 14.7 mg/mL in Morpheus G12 (0.1 M Buffer Sytem 3 pH 8.5, 0.1 M Carboxylic acids, 37.5% v/v Precipitant Mix 4) at 18°C after 2 months. Crystals were cryo-cooled in liquid nitrogen, then diffraction data collected at Diamond Light Source on beamline I04 (wavelength 0.9537 Å) which auto-processed with xia2-dials^41^ to 2.20 Å. Molecular replacement (MR) was performed with the CCP4^40^ implementation of PHASER^42^ using the constant and variable domains of a mouse antibody (PDB 5KVG) trimmed to remove CDRs as separate search models. Searches for PfRIPR domains EGF6 or EGF7 using their AlphaFold^44^ predictions were unsuccessful (using domains isolated from the AlphaFold prediction of PfRIPR^tail^) and thus PfRIPR was built *de novo* by iterative model building in COOT (0.9.3)^43^ and BUSTER (2.10.4)^48^, by first building the CDRs of RP.012, then backbone tracing PfRIPR with a poly-Ala chain followed by sequence identification and building. In this way, we built PfRIPR residues 781-787, 798-843 and 847-853. The unit cell contains two copies of the RP.012-PfRIPR complex. In both copies, EGF6 and EGF7 are incomplete, thus the AlphaFold predictions of each domain and their conserved disulphide bonds were used as guides in the later stages of model building. These AlphaFold predictions align well with the resolved regions of PfRIPR with backbone RMSDs of 0.47 Å at EGF6 (over 25 pruned atom pairs, 0.69 Å over all 26 pairs) and 0.95 Å at EGF7 respectively (over 27 pruned atom pairs, 1.23 Å across all 33 pairs). The centre of mass of each of these aligned AlphaFold predicted domains were used to measure the angle between domains using ChimeraX^45^ during structure analysis.

A complex of the single chain variable fragment (scFv) of MAD8-739 and PfRIPR EGF5-6 was isolated by gel filtration using an S75 Increase 10/300 in HBS pH 7.5. We obtained crystals using sitting drop vapour diffusion as above using 9.7 mg/mL protein complex in ProPlex B9 (0.1 M sodium cacodylate pH 6.0, 15% w/v PEG 4000) at 18°C after 2 months. Crystals were cryo-protected using the reservoir solution supplemented with 25% v/v glycerol then cryo-cooled in liquid nitrogen. Diffraction data were collected at Diamond Light Source on beamline I24 (wavelength 0.6199 Å). After auto-processing in xia2-dials, unmerged data were then further processed using AIMLESS^46^ in CCP4 to 2.8 Å. MR was performed by PHASER using a variable domain with truncated CDRs as a search model. As with RP.012, PfRIPR (residues 719-816) was built by iterative refinement. The unit cell contains two copies of the MAD8.739-PfRIPR complex.

The crystal structure of MAD8-66 and PfRIPR EGF9-CTD was obtained after reconstituting a complex comprising the Fab fragment of MAD8-66, PfRIPR EGF9-CTD, PfPTRAMP-PfCSS, and nanobodies D2 and H8^3^, isolated by gel filtration using a SRT-10C SEC-300 HPLC column (Sepax Technologies) in 20 mM HEPES pH 7.5, 50 mM NaCl. Crystals were obtained using sitting drop vapour diffusion as above using 9.9 mg/mL protein complex in JCSG+ B5 (0.1 M sodium cacodylate pH 6.5, 40% w/v MPD, 5% w/v PEG 8000) at 18°C. These were cryo-protected in reservoir solution plus 25% v/v glycerol, and cryo-cooled in liquid nitrogen. Diffraction data were collected at Diamond Light Source on beamline I04 (wavelength 0.9357 Å), and auto-processed using xia2-3dii to 2.25 Å. MR was performed using PHASER with antibody Fab fragment heavy chain (PDB 7BEH) and light chains (PDB 6DWA), trimmed to remove their CDRs, and the AlphaFold prediction of PfRIPR CTD as search models. The remainder of PfRIPR (EGF9 and 10, obtaining a final model comprising 907-916, 922-1021 and 1025-1082 of PfRIPR) was built by iterative refinement as above, with later refinement cycles performed in PHENIX (1.21.2)^47^. No PfPTRAMP-PfCSS nor nanobodies D2 and H8 were observed indicating that these had dissociated prior to crystal formation. The unit cell contains a single copy of the MAD8.66-PfRIPR complex.

#### Cryo-EM data collection

A complex of PfRIPR^67L^ (PfRIPR with a designed disulphide bond between domains EGF6 and 7 as described above), PfCyRPA, PfRH5 and the Fab fragments of Cy.007 and MAD8-2 was isolated by gel filtration using a S200 Increase 10/300 in HBS pH 7.5. Fractions containing the intact complex were concentrated using an 100K Amicon Ultra centrifugal unit at 6,000 g and 4°C. Grids were prepared with a FEI Vitrobot Mark IV (Thermo Fisher) by applying 3 µL of complex at 0.25 mg/mL to C-Flat 1.2/1.3 grids (Electron Microscopy Sciences) which had been glow-discharged for 60 s at 15 mA. Samples were incubated for 5 s at 4°C and 100% humidity, then grids blotted for 4 s and plunge frozen into liquid ethane.

Data were collected using an FEI Titan Krios operating at 300 kV equipped with a Gatan K3 direct electron detector and a Gatan BioQuantum 20 eV energy filter. Data collection was automated using fast acquisition mode in EPU (ThermoFisher). Movies were acquired at x58,149 magnification with a calibrated pixel size of 0.83 Å per pixel, a total dose of 39.7 electrons per Å^2^ over 2.39 s with 40 frames, and a defocus range of -2.6 to -1.2 µm in 0.2 µm increments. A total of 28,491 movies were collected.

#### Cryo-EM image processing

Image processing was performed using cryoSPARC v4 and RELION 5.0.0 and is illustrated in Figure S4B. Movies were motion corrected with patch motion in cryoSPARC, then the contrast transfer function (CTF) of micrographs estimated using patch CTF. Micrographs were curated to remove those with a relative ice thickness greater than 1.2 or an estimated CTF fit resolution greater than 6.

A subset of 1,276 micrographs was blob picked using an elliptical blob with a particle diameter of 50-200 Å. 500,999 particles were extracted (at 1.66 Å/pixel) and 2D classified for multiple iterations to yield templates for template picking. 472,845 particles from this template picking were extracted and 2D classified as before. This yielded higher quality 2D classes that were used to template pick the full dataset of 26,624 micrographs yielding 10,384,775 particles. These were iteratively 2D classified to separate and discard poor quality particles, obtaining a final stack of 4,134,414 particles. These were subjected to *ab initio* reconstruction then heterogenous refinement with a junk decoy volume. These volumes and 2D classes indicated that the particle stack was heterogeneous, containing the PfRCR-Cy.007-MAD8-2 complex, a PfRIPR-PfCyRPA-Cy.007 complex, and a Fab fragment bound a small antigen. This antigen had a distinct arch shape identical to the tail of PfRIPR seen in the PfRCR-Cy.007-MAD8-2 complex, thus we concluded that these particles were of MAD8-2 bound to PfRIPR. Before proceeding, micrographs were repicked using TOPAZ^57^, particles extracted and duplicates removed, obtaining a combined stack of 7,476,072 particles.

We first sought to resolve the intact PfRCR-Cy.007-MAD8-2 complex. Following iterative 2D classification of our combined particles, we manually separated particles into stacks for each species in our dataset using their 2D classes. 969,432 particles corresponding to PfRCR-Cy.007-MAD8-2 were homogenously, then non-uniform refined^58^ to obtain a Nyquist-limited volume at 3.4 Å resolution. Then, to resolve MAD8-2, we performed 3D classification into 10 classes. Three of these classes showed clear density for the tail of PfRIPR and MAD8-2. We combined the 257,180 particles from these three classes, then performed reference-based motion correction and re-extracted them at their full pixel size (0.83 Å/pixel). Non-uniform refinement of these ‘polished’ particles led to a volume with a global 3.0 Å resolution. While PfCyRPA, Cy.007, and the core of PfRIPR were well resolved in this map (with a local resolution of 2.5-3.0 Å), the tail of PfRIPR and MAD8-2 were not (∼6.5 Å local resolution). To improve density for PfRIPR tail, we re-extracted our particles re-centred on to PfRIPR and MAD8-2, then performed local refinement using a mask around these components, yielding an improved map at 2.8 Å resolution. This map showed clear improved density for EGF5 and the N-terminus of PfRIPR and was used for model building, detailed later.

We next sought to resolve the MAD8-2-PfRIPR complex. To do this, we re-extracted our particle stack of MAD8-2 bound to a small fragment of PfRIPR (669,915 particles at 0.83 Å/pixel), then performed *ab initio* reconstruction and heterogenous refinement to isolate a clean stack of 377,702 particles. These were subject to homogeneous refinement, then non-uniform refinement to obtain a volume at 3.3 Å resolution. We next performed local refinement using a mask around PfRIPR and the variable domain of MAD8-2, yielding a volume with a reported 3.4 Å resolution. However, this volume was visibly anisotropic due to limited views. Noting that 2D classification showed that our PfRCR-Cy.007-MAD8-2 particle stack had different views of MAD8-2, we sought to combine our particle stacks to improve the volume. To achieve this, we re-centred our PfRCR-Cy.007-MAD8-2 volume on to MAD8-2 using the volume align tool in cryoSPARC, then re-extracted and combined these with our 377,702 particles of PfRIPR-MAD8-2 alone (Figure S4B). These combined particles were 2D classified to further clean the particle stack to 318,577 particles, which were subject to reference-based motion correction, then non-uniform refined to 3.1 Å. Given the low molecular weight of this Fab-PfRIPR complex, we exported these particles into RELION (5.0.0) to make use of BLUSH regularisation^50^, an algorithm which achieves better quality reconstructions for small macromolecules. In RELION, we first performed 3D classification without alignment (4 classes, T value = 4, masked to MAD8-2 variable domain), then took the best volume (191,422 particles) for 3D refinement with BLUSH. This yielded a 3.5 Å volume, which had visible separation of beta strands in both MAD8-2 variable domain and PfRIPR, and was of better quality than that obtained in cryoSPARC despite the nominally lower reported resolution. We used this RELION 3.5 Å volume to model the MAD8-2-PfRIPR complex as detailed below.

#### Cryo-EM model building

To build a structure of PfRIPR-PfCyRPA-Cy.007, we first focused on PfRIPR. The published structure of PfRIPR core (PDB entry 8CDD) was docked into our PfRIPR-focused volume using ChimeraX, then the model extended iteratively through cycles of building in COOT (0.9.3) and real-space refinement with secondary structure restraints in PHENIX (1.21.2). The AlphaFold prediction of PfRIPR N-EGF5 was used to guide model building. This final model of PfRIPR comprises residues 25-769 (excluding disordered loops at 49-51, 104-109, 124-137, 276-279 and 478-556), extending the known structure of PfRIPR by 9 and 53 residues at the N and C-termini respectively. Next, we docked the published model of Cy.007-bound PfCyRPA (PDB entry 7PHV) into our volume and refined the model using PHENIX as above. As density corresponding to Cy.007 was limited in our map, we did not include it in our final model, which contains PfRIPR and PfCyRPA only. This model shows some new contacts between PfRIPR and PfCyRPA not present in the published structure of PfRCR (PDB entry 8CDD).

To build a model of MAD8-2 and PfRIPR, we predicted the structure of PfRIPR EGF6 and MAD8-2 variable domain using AlphaFold3^44^ with 20 different seeds, guided by our prior competition epitope binning and structural analyses. We ranked these predictions using their ipTM scores, then docked the highest-ranking model (ipTM of 0.91, pTM of 0.89), plus an AlphaFold prediction of the Fab constant domain of MAD8-2 into our 3.5 Å volume of PfRIPR-MAD8-2 using ChimeraX. We relaxed this model into the cryo-EM volume using ISOLDE^49^, then performed real-space refinement using secondary structure restraints in PHENIX (1.21.2).

#### Modelling the PfPCRCR complex

To generate a composite model of the PfPCRCR complex, we aligned structures of PfRCR determined previously (PDB entry 8CDD), PfRIPR N-EGF5 (from our PfRIPR-PfCyRPA-Cy.007 structure), PfRIPR EGF5-6 (from our MAD8-739-bound PfRIPR), AlphaFold3 models of PfRIPR tail (residues 717-1086) and the PfPTRAMP-PfCSS-PfRIPR complex (PfPTRAMP residues 42-352, PfCSS residues 21-290, and PfRIPR residues 717-1086). Alignments were restricted to the core of PfRIPR (N-EGF4), PfRIPR EGF5, PfRIPR EGF6 and PfRIPR CTD respectively where required to assemble this composite model. Later, the crystal structure of RP.012-bound PfRIPR was used to visualise the effect of RP.012 binding on the conformation of the PfPCRCR complex, using EGF7 to drive the alignment. AlphaFold3 predictions were performed using the AlphaFold Server. These predictions were coloured by pLDDT and Predicted Alignment Error (PAE) plots visualised using PAE Viewer^51^.

#### Quantification and statistical analysis

Details of statistical calculations for each experiment, including sample size, statistical test, average and measure of spread, can be found in the relevant figure legends.

---

**Figure S1:**
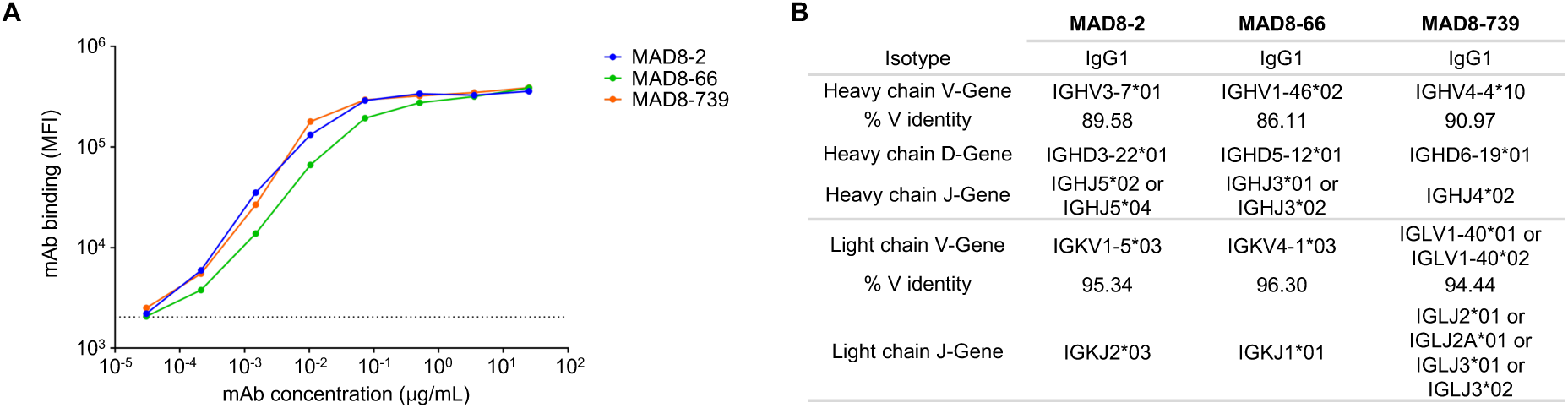
PfRIPR-specific human mAbs isolated from malaria-exposed individuals, related to Figure 1. (A) Binding of mAbs MAD8-2, MAD8-66 and MAD8-739 to PfRIPR-coated beads. (B) Gene and isotype information for MAD8-2, MAD8-66 and MAD8-739. Gene information was obtained from the International Immunogenetics Information System (IMGT) database.

**Figure S2:**
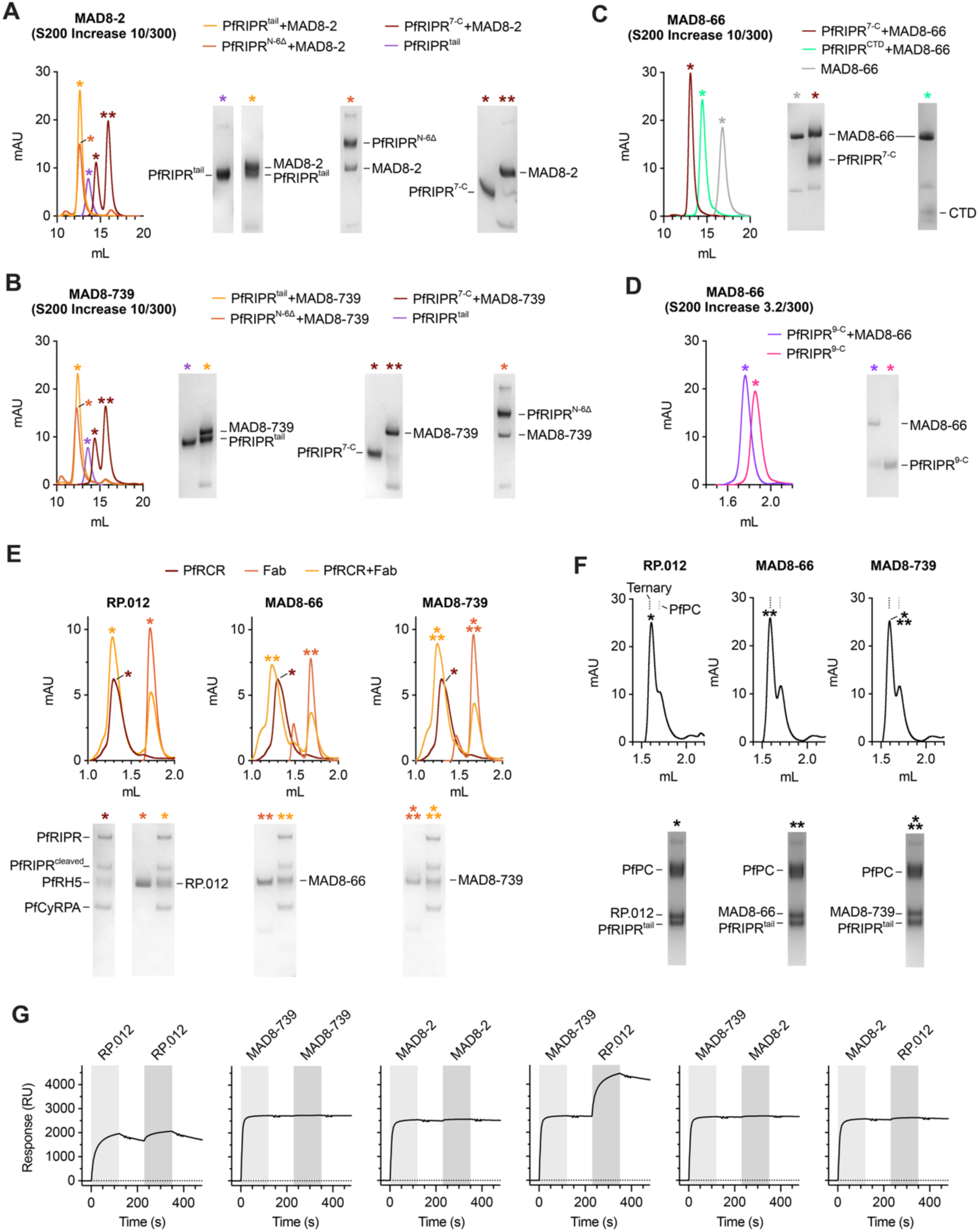
Epitope binning and competition analysis of PfRIPR antibodies, related to Figure 1. (A-D) SEC traces of MAD8-2, MAD8-66 and MAD8-739 Fab fragments with and without various fragments of PfRIPR to identify their epitopes, as summarised in Figure 1F; Fab fragment (grey), PfRIPR^tail^ (purple), PfRIPR^tail^+Fab (orange), PfRIPR^N-6Δ^+Fab (red), PfRIPR^7-C^+Fab (brown), PfRIPR^CTD^+Fab (light green), PfRIPR^9-C^ (pink), PfRIPR^9-C^+Fab (purple). For each analysis, peak fractions as identified by coloured asterisks were analysed by SDS-PAGE to verify co-elution of components. (E) SEC traces of RP.012, MAD8-66 and MAD8-739 Fab fragments (red), PfRCR (brown) and their mixture (orange), illustrating co-elution of all Fabs with the intact PfRCR complex. Peak fractions, identified by the coloured asterisks, were analysed by SDS-PAGE to confirm co-elution of components. The PfRCR trace and SDS-PAGE gel lane is replicated from Figure 1G for comparisons in the presence of each Fab fragment. (F) SEC traces of RP.012, MAD8-66 or MAD8-739 Fab fragments after incubation with PfPTRAMP-PfCSS and PfRIPR tail. The dotted lines indicate the elution volume of the ternary complex peak observed for MAD8-2 (black) and PfPTRAMP-PfCSS (grey) as shown in Figure 1H, illustrating that each antibody forms a ternary complex that elutes similarly. The fraction at this location was analysed by SDS-PAGE to verify co-elution of all three components. (G) Representative raw SPR traces showing sequential injection of Fab fragments of EGF5-8 binding antibodies over PfRIPR immobilised on an SPR chip, demonstrating binding competition; n=3, 1 shown.

**Figure S3:**
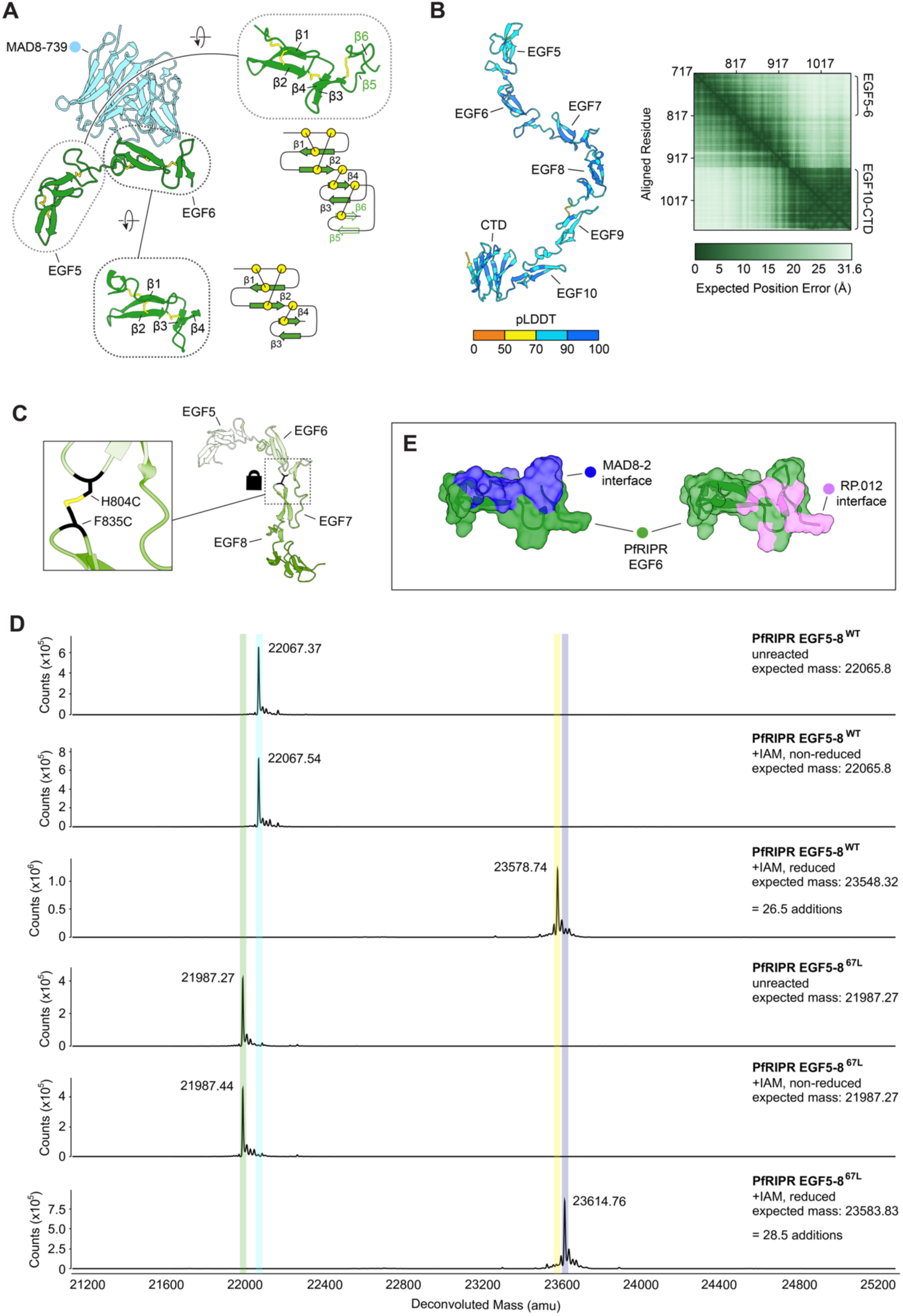
Structural analysis of PfRIPR and design of a lock between PfRIPR EGF6-7, related to Figure 2. (A) Structure of MAD8-739 (light blue) and PfRIPR EGF-like domains 5-6 (green) in cartoon representation with disulphide bonds shown as yellow sticks. Each domain is expanded and shown inset, to display these disulphide bonds more clearly, with the major (β1-β2) and minor sheets (β3-β4) of the EGF-like fold labelled. EGF-like domain 5 contains an additional disulphide bond, and an extra structural element like an additional beta sheet (β5-β6), like a laminin fold^27^. A schematic representation of each EGF-like domain is shown adjacent to each inset. (B) AlphaFold prediction of PfRIPR^tail^ in cartoon representation coloured by pLDDT (left) and its PAE matrix (right) with domains of interest labelled. (C) Design of a cysteine-locked PfRIPR through introduction of a disulphide bond between EGF6 and 7 at H804 and F835 (as shown inset). (D) Intact mass analysis of wild-type (EGF5-8^WT^) or EGF6-7 cysteine-locked PfRIPR (EGF5-8^67L^), either unreacted, or following reaction with iodoacetamide (IAM) under denaturing conditions both with and without reducing agent. The expected and detected deconvoluted masses are shown with bars highlighting masses observed for clarity. PfRIPR EGF5-8^WT^ and PfRIPR EGF5-8^67L^ match their expected unlabelled masses in the absence of reducing agent (green and blue bars, respectively), indicating that they each contain no free cysteine residues. Under reducing conditions, EGF5-8^WT^ is labelled with 26.5 carbamidomethyl groups indicating that it contains 13 disulphide bonds (yellow bar), and EGF5-8^67L^ is labelled with 28.5 carbamidomethyl groups indicating that it contains 14 disulphide bonds (13 native and 1 introduced, purple bar). (E) EGF6 in cartoon and surface representation with the interaction interface of MAD8-2 (dark blue) and RP.012 (pink) highlighted, showing overlap of interfaces.

**Figure S4:**
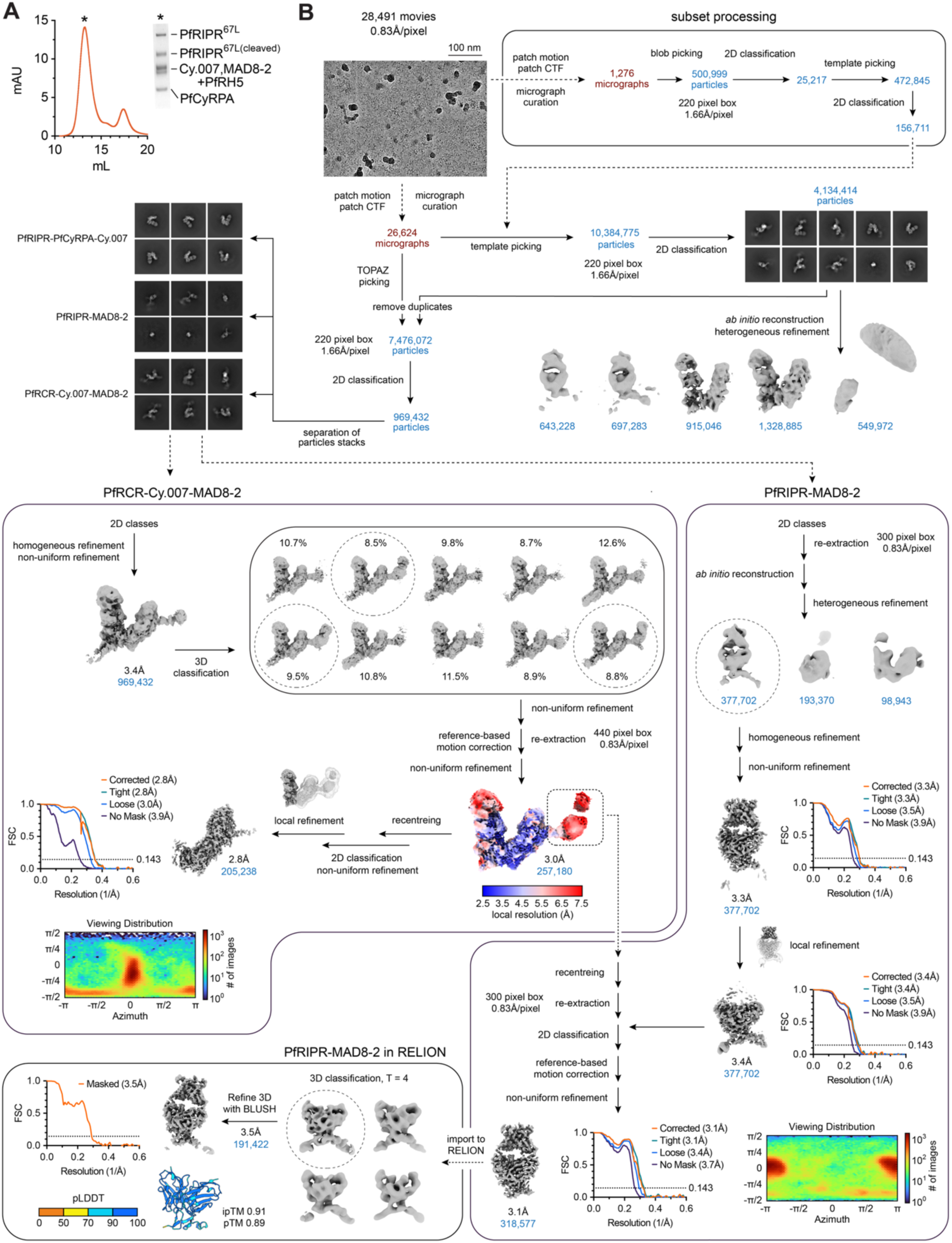
Structure determination of MAD8-2-bound PfRIPR, related to Figure 2 and STAR Methods. (A) Size exclusion chromatography trace showing isolation of a MAD8-2- and Cy.007-bound PfRCR complex with the peak fraction analysed by SDS-PAGE shown inset. (B) Processing pipeline of single particle cryo-EM data. Early processing indicated that the dataset contained two particle stacks of interest, with one of the MAD8-2-bound PfRCR-Cy.007 complex and one of MAD8-2-bound PfRIPR. These were separated and initially processed separately but partially recombined later following particle recentring to aid MAD8-2-bound PfRIPR volume refinement. The predicted structure of this complex was predicted by AlphaFold3, shown in cartoon representation coloured by pLDDT.

**Figure S5:**
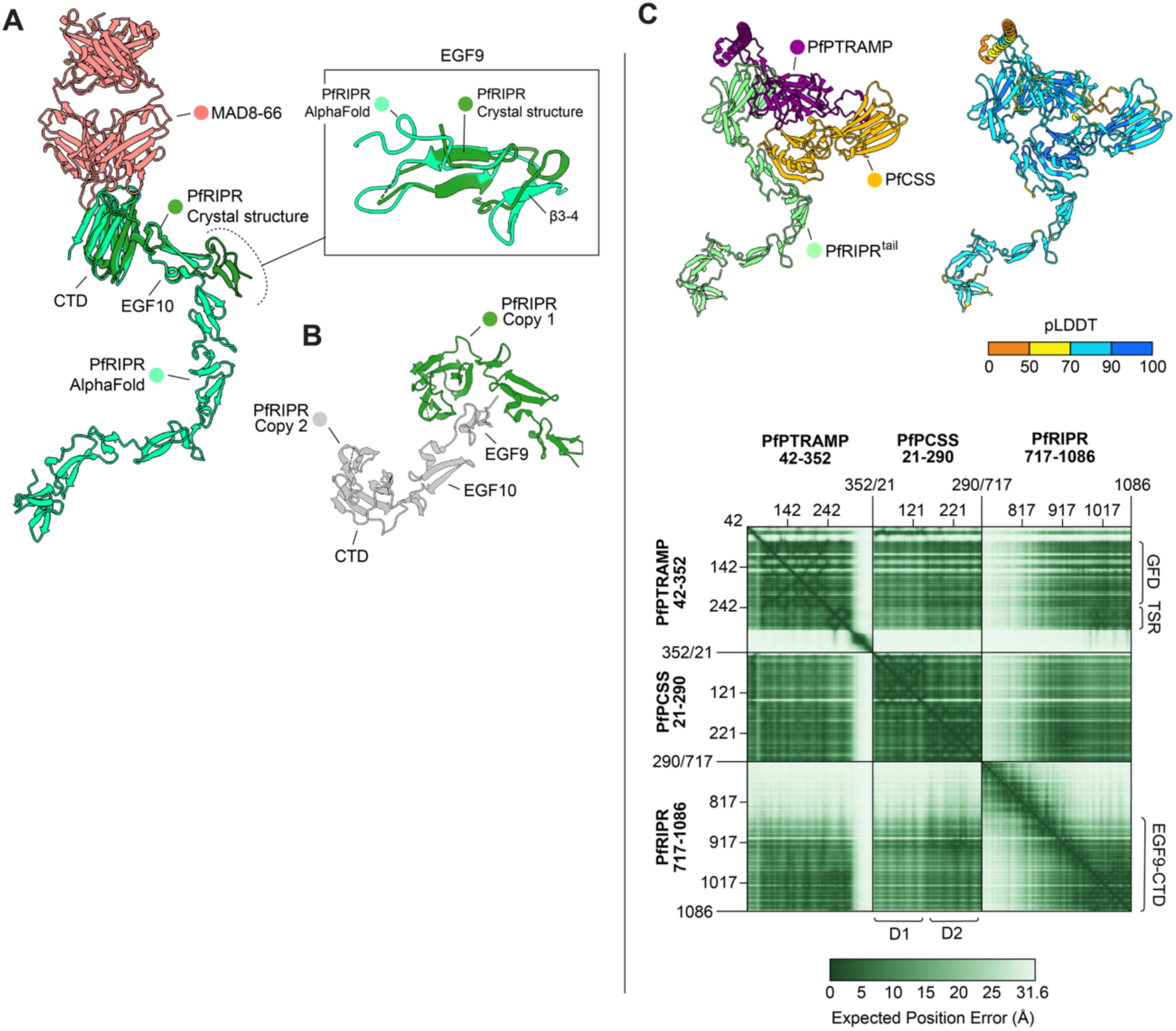
Analysis of PfRIPR-antibody structures, related to Figure 3. (A) Alignment of the crystal structure of MAD8-66 (pink) and PfRIPR EGF9-CTD (green) and the AlphaFold prediction of PfRIPR^tail^ (light green), illustrating different placements of EGF9 and a different conformation in β3-β4 of EGF9 (shown inset). (B) Symmetry mates of PfRIPR EGF9-CTD (green and grey) illustrating that EGF9 forms crystal contacts with the CTD of its symmetry mate, likely inducing conformational changes in EGF9 (Figures 3B and S5A). (C) AlphaFold3 prediction of PfPTRAMP-PfCSS-PfRIPR^tail^ in cartoon representation coloured by chain (left) and by pLDDT (right), and its PAE matrix (below) with domains of interest labelled.

**Table S1:**
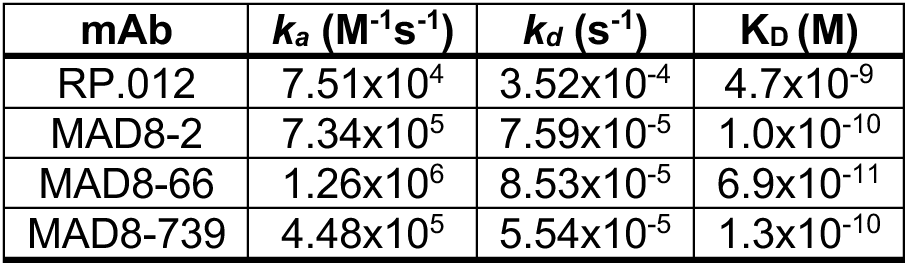
Kinetic and affinity parameters for mAb binding to PfRIPR.

**Table S2:** X-ray crystallography data collection and refinement statistics.

|  | **RP.012-bound PfRIPR**<br>PDB ID 9TUW | **MAD8-66-bound PfRIPR**<br>PDB ID 9TUX | **MAD8-739-bound PfRIPR**<br>PDB ID 9TUY |
| --- | --- | --- | --- |
| **Data Collection** |  |  |  |
| Space group | P 1 | P 21 21 21 | P 3 1 |
| Cell dimensions |  |  |  |
| a, b, c (Å) | 50.16, 50.98, 109.78 | 52.79, 63.24, 250.73 | 89.64, 89.64, 141.34 |
| $\alpha$, $\beta$, $\gamma$ (°) | 76.82, 83.26, 76.50 | 90.00, 90.00, 90.00 | 90.00, 90.00, 120.00 |
| Wavelength | 0.9537 Å | 0.9537 Å | 0.6199 Å |
| Resolution (Å) | 48.67-2.20<br>(2.23-2.20) | 63.25-2.25<br>(2.29-2.25) | 42.72-2.80<br>(2.95-2.80) |
| Total Observations | 186049 (9015) | 549427 (28167) | 335117 (49712) |
| Total Unique | 51504 (2453) | 41042 (2058) | 31298 (4549) |
| $R_{\text{merge}}$ (%) | 10.1 (132.7) | 29.4 (410.8) | 32.6 (379.2) |
| $R_{\text{meas}}$ (%) | 11.9 (155.5) | 30.6 (426.6) | 34.3 (397.8) |
| $R_{\text{pim}}$ (%) | 6.2 (80.6) | 8.3 (41.5) | 10.5 (120.2) |
| $CC_{1/2}$ | 0.995 (0.317) | 0.996 (0.415) | 0.993 (0.810) |
| $I/\sigma(I)$ | 11.1 (0.8) | 8.3 (0.7) | 5.6 (1.1) |
| Completeness (%) | 98.3 (94.3) | 100.0 (100.0) | 100.0 (100.0) |
| Multiplicity | 3.6 (3.7) | 13.4 (13.7) | 10.7 (10.9) |
| Wilson B factor | 53.41 | 47.08 | 68.7 |
| **Refinement** |  |  |  |
| Reflections | 51362 | 40916 | 31142 |
| Rwork / Rfree (%) | 24.1/25.6 | 23.0/26.2 | 22.8/25.5 |
| Average B factor |  |  |  |
| Protein | 65.67 | 56.58 | 85.10 |
| Water | 55.29 | 54.16 | 71.16 |
| Number of residues |  |  |  |
| Protein | 959 | 605 | 674 |
| Water | 157 | 93 | 63 |
| RMSDs |  |  |  |
| Bond lengths (Å) | 0.010 | 0.003 | 0.010 |
| Bond angles (°) | 1.40 | 0.58 | 1.42 |
| Ramachandran plot |  |  |  |
| Favored (%) | 98.0 | 98.0 | 94.4 |
| Allowed (%) | 2.0 | 2.0 | 5.6 |
| Outliers (%) | 0 | 0 | 0 |
| Clashscore | 5.36 | 2.80 | 3.00 |
| MolProbity score | 1.82 | 1.07 | 1.72 |

**Table S3:** Interaction contacts between RP.012 and PfRIPR.

| PfRIPR |  |  |  | RP.012 |  |  |  | Interaction |
| --- | --- | --- | --- | --- | --- | --- | --- | --- |
| Chain | Residue | Group | Location | Chain | Residue | Group | Location |  |
| D | E800 | Side chain | EGF6 | F | S87 | Side chain | Kappa FR3 | H-bond |
| D | E803 | Side chain | EGF6 | F | S70 | Side chain | CDR L2 | H-bond |
| D | E803 | Backbone | EGF6 | F | T51 | Side chain | CDR L1 | H-bond |
| D | Y806 | Side chain | EGF6 | F | R113 | Side chain | CDR L3 | Cation-pi |
| D | R807 | Backbone | EGF6 | F | N48 | Side chain | CDR L1 | H-bond |
| D | N816 | Side chain | EGF6 | F | S112 | Side chain | CDR L3 | H-bond |
| D | D817 | Backbone | EGF6 | F | S112 | Backbone | CDR L3 | H-bond |
|  | D817 | Side chain | EGF7 | F | Y114 | Backbone | CDR L3 | H-bond |
|  |  |  |  | F | L66 | Side chain | Kappa FR2 | Hydrophobic |
|  |  |  |  | F | Y69 | Side chain | Kappa FR2 | Hydrophobic |
|  |  |  |  | F | Y111 | Side chain | CDR L3 | Hydrophobic |
|  |  |  |  | F | F116 | Side chain | CDR L3 | Hydrophobic |
| D | Y818 | Side chain | EGF7 |  |  |  |  | Hydrophobic |
| D | D821 | Side chain | EGF7 | E | R70 | Side chain | Heavy FR2 | Salt bridge |
|  | D821 | Side chain | EGF7 | E | K79 | Side chain | Heavy FR3 | Salt bridge |
|  | D821 | Side chain | EGF7 | F | Y114 | Side chain | Heavy FR3 | H-bond |
|  | D821 | Side chain | EGF7 | E | R70 | Side chain | Heavy FR2 | H-bond |
| D | I832 | Side chain | EGF7 |  |  |  |  | Hydrophobic |
| D | N834 | Backbone | EGF7 | E | Y52 | Side chain | CDR H1 | H-bond |
| D | N834 | Side chain | EGF7 | E | Y52 | Side chain | CDR H1 | H-bond |
| D | F835 | Backbone | EGF7 | E | L120 | Backbone | CDR H3 | H-bond |
|  | F835 | Side chain | EGF7 | E | D123 | Side chain | CDR H3 | Anion-pi |
| D | F835 | Side chain | EGF7 |  |  |  |  | Hydrophobic |
| D | P837 | Side chain | EGF7 |  |  |  |  | Hydrophobic |
|  |  |  |  | E | W53 | Side chain | CDR H1 | Hydrophobic |
|  |  |  |  | E | L120 | Side chain | CDR H3 | Hydrophobic |

**Table S4:**
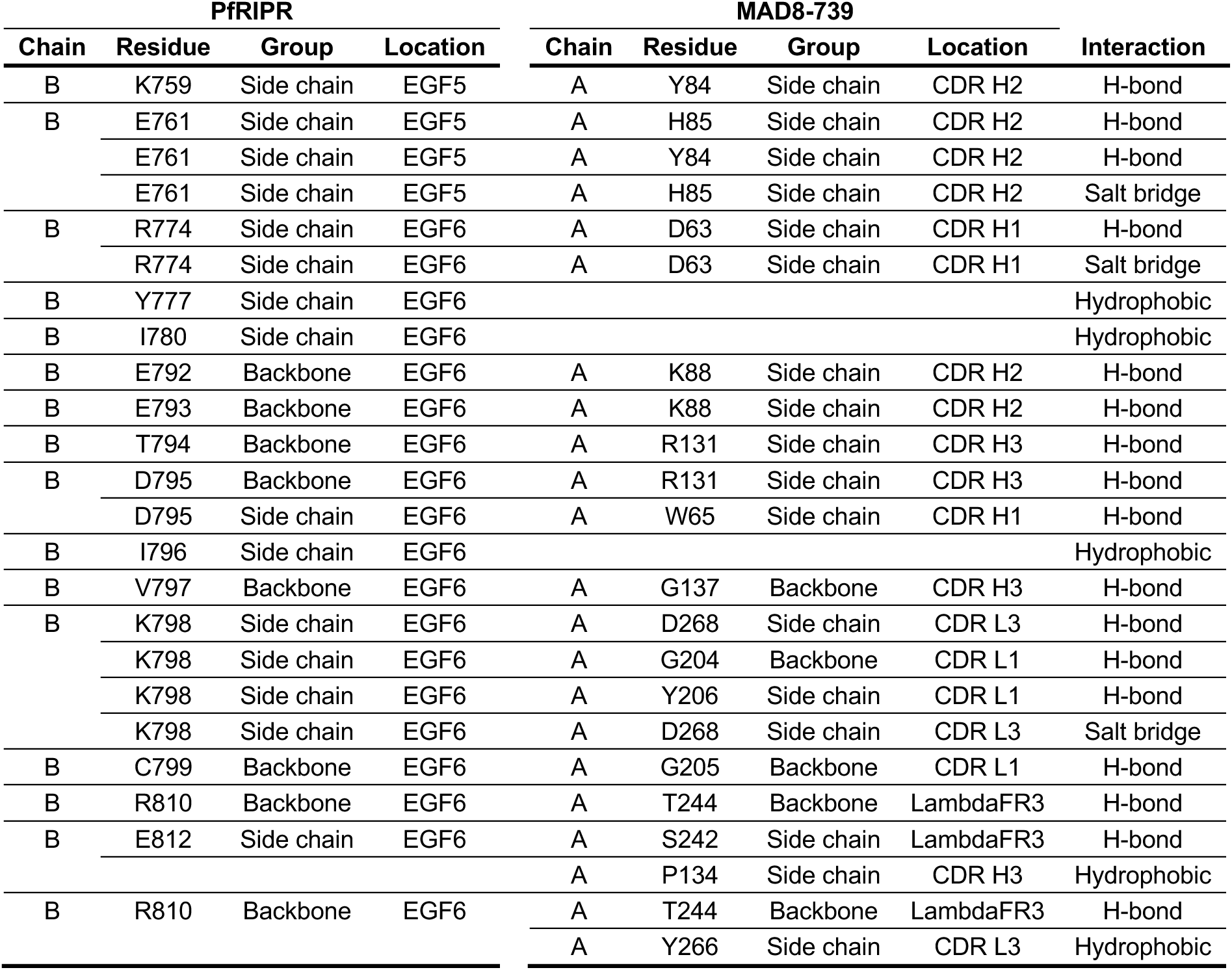
Interaction contacts between MAD8-739 and PfRIPR.

| PfRIPR |  |  |  | MAD8-739 |  |  |  | Interaction |
| --- | --- | --- | --- | --- | --- | --- | --- | --- |
| Chain | Residue | Group | Location | Chain | Residue | Group | Location |  |
| B | K759 | Side chain | EGF5 | A | Y84 | Side chain | CDR H2 | H-bond |
| B | E761 | Side chain | EGF5 | A | H85 | Side chain | CDR H2 | H-bond |
|  | E761 | Side chain | EGF5 | A | Y84 | Side chain | CDR H2 | H-bond |
|  | E761 | Side chain | EGF5 | A | H85 | Side chain | CDR H2 | Salt bridge |
| B | R774 | Side chain | EGF6 | A | D63 | Side chain | CDR H1 | H-bond |
|  | R774 | Side chain | EGF6 | A | D63 | Side chain | CDR H1 | Salt bridge |
| B | Y777 | Side chain | EGF6 |  |  |  |  | Hydrophobic |
| B | I780 | Side chain | EGF6 |  |  |  |  | Hydrophobic |
| B | E792 | Backbone | EGF6 | A | K88 | Side chain | CDR H2 | H-bond |
| B | E793 | Backbone | EGF6 | A | K88 | Side chain | CDR H2 | H-bond |
| B | T794 | Backbone | EGF6 | A | R131 | Side chain | CDR H3 | H-bond |
| B | D795 | Backbone | EGF6 | A | R131 | Side chain | CDR H3 | H-bond |
|  | D795 | Side chain | EGF6 | A | W65 | Side chain | CDR H1 | H-bond |
| B | I796 | Side chain | EGF6 |  |  |  |  | Hydrophobic |
| B | V797 | Backbone | EGF6 | A | G137 | Backbone | CDR H3 | H-bond |
| B | K798 | Side chain | EGF6 | A | D268 | Side chain | CDR L3 | H-bond |
|  | K798 | Side chain | EGF6 | A | G204 | Backbone | CDR L1 | H-bond |
|  | K798 | Side chain | EGF6 | A | Y206 | Side chain | CDR L1 | H-bond |
|  | K798 | Side chain | EGF6 | A | D268 | Side chain | CDR L3 | Salt bridge |
| B | C799 | Backbone | EGF6 | A | G205 | Backbone | CDR L1 | H-bond |
| B | R810 | Backbone | EGF6 | A | T244 | Backbone | LambdaFR3 | H-bond |
| B | E812 | Side chain | EGF6 | A | S242 | Side chain | LambdaFR3 | H-bond |
|  |  |  |  | A | P134 | Side chain | CDR H3 | Hydrophobic |
| B | R810 | Backbone | EGF6 | A | T244 | Backbone | LambdaFR3 | H-bond |
|  |  |  |  | A | Y266 | Side chain | CDR L3 | Hydrophobic |

**Table S5:** Kinetic and affinity parameters for mAb binding to PfRIPR^67L^.

| **mAb** | **$k_a$ (M<sup>-1</sup>s<sup>-1</sup>)** | **$k_d$ (s<sup>-1</sup>)** | **$K_d$ (M)** |
| --- | --- | --- | --- |
| RP.012 | n/a | n/a | n/a |
| MAD8-2 | 8.61x10<sup>5</sup> | 9.68x10<sup>-5</sup> | 1.1x10<sup>-10</sup> |

**Table S6:**
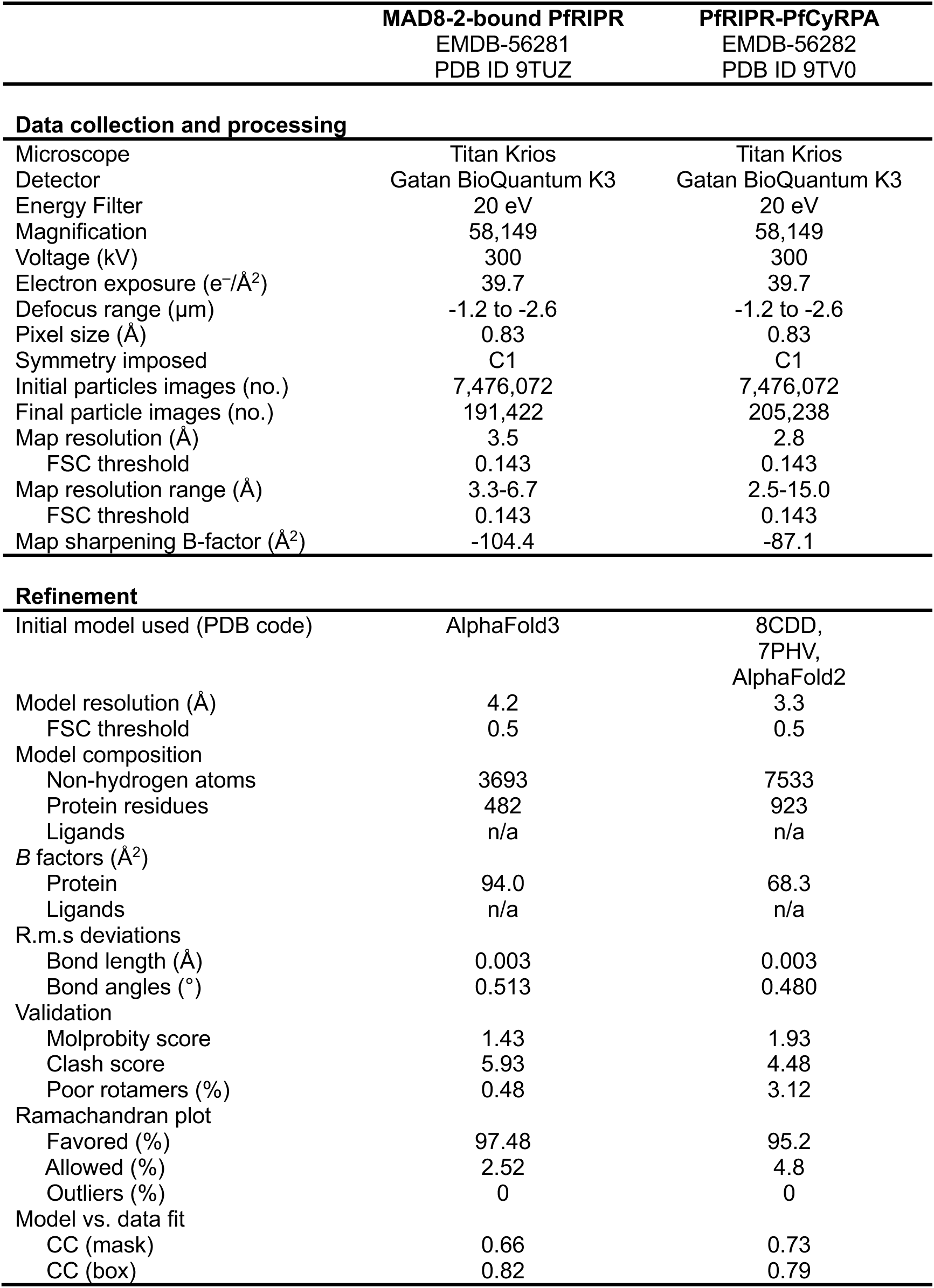
Cryo-EM data collection and refinement statistics.

**Table S7:** Interaction contacts between MAD8-66 and PfRIPR.

| PfRIPR |  |  |  | MAD8-66 |  |  |  | Interaction |
| --- | --- | --- | --- | --- | --- | --- | --- | --- |
| Chain | Residue | Group | Location | Chain | Residue | Group | Location |  |
| C | E983 | Side chain | EGF10-CTD | B | N54 | Side chain | CDR H2 | H-bond |
| C | Y985 | Side chain | EGF10-CTD | B | D28 | Side chain | CDR H1 | H-bond |
| C | I1000 | Side chain | CTD |  |  |  |  | Hydrophobic |
| C | A1003 | Side chain | CTD |  |  |  |  | Hydrophobic |
| C | N1045 | Backbone | CTD | B | K52 | Side chain | CDR H2 | H-bond |
| C | E1046 | Side chain | CTD | B | N57 | Side chain | CDR H2 | H-bond |
|  | E1046 | Side chain | CTD | B | K52 | Side chain | CDR H2 | Salt bridge |
| C | I1047 | Side chain | CTD |  |  |  |  | Hydrophobic |
| C | F1048 | Side chain | CTD |  |  |  |  | Hydrophobic |
| C | T1073 | Backbone | CTD | A | S99 | Side chain | CDR L3 | H-bond |
|  | T1073 | Backbone | CTD | A | A100 | Backbone | CDR L3 | H-bond |
| C | C1074 | Side chain | CTD | A | S98 | Backbone | CDR L3 | H-bond |
| C | I1075 | Side chain | CTD |  |  |  |  | Hydrophobic |
| C | E1077 | Backbone | CTD | B | D104 | Side chain | CDR H3 | H-bond |
|  | E1077 | Side chain | CTD | B | F33 | Backbone | CDR H1 | H-bond |
|  | E1077 | Side chain | CTD | B | H35 | Side chain | Heavy FR2 | Salt bridge |
| C | N1078 | Backbone | CTD | B | D104 | Side chain | CDR H3 | H-bond |
|  | N1078 | Side chain | CTD | B | D104 | Side chain | CDR H3 | H-bond |
|  | N1078 | Side chain | CTD | B | H100 | Backbone | CDR H3 | H-bond |
| C | F1080 | Side chain | CTD | A | N34 | Side chain | CDR L1 | Polar-pi |
|  |  |  |  | A | F31 | Side chain | CDR L1 | Hydrophobic |
|  |  |  |  | A | A33 | Side chain | CDR L1 | Hydrophobic |
|  |  |  |  | A | Y38 | Side chain | CDR L1 | Hydrophobic |
|  |  |  |  | A | W56 | Side chain | CDR L2 | Hydrophobic |
|  |  |  |  | A | A100 | Side chain | CDR L3 | Hydrophobic |
|  |  |  |  | A | W102 | Side chain | CDR L3 | Hydrophobic |
|  |  |  |  | B | F33 | Side chain | CDR H1 | Hydrophobic |
|  |  |  |  | B | L50 | Side chain | Heavy FR2 | Hydrophobic |
|  |  |  |  | B | A59 | Side chain | Heavy FR3 | Hydrophobic |
|  |  |  |  | B | V99 | Side chain | CDR H3 | Hydrophobic |

